# Cryo-EM of a viral RNA and RNA-protein complex reveals how structural dynamics and novel tRNA mimicry combine to hijack host machinery

**DOI:** 10.1101/2020.09.18.302638

**Authors:** Steve L. Bonilla, Madeline E. Sherlock, Andrea MacFadden, Jeffrey S. Kieft

## Abstract

Viruses require multifunctional structured RNAs to hijack their host’s biochemistry, but their mechanisms can be obscured by the difficulty of solving conformationally dynamic RNA structures. Using cryo-EM, we visualized the structure of the mysterious viral tRNA-like structure (TLS) from brome mosaic virus (BMV), which affects replication, translation, and genome encapsidation. Structures in isolation and bound to tyrosyl-tRNA synthetase (TyrRS) show that this ∼55 kDa purported tRNA mimic undergoes large conformational rearrangements to bind TyrRS in a form that differs dramatically from tRNA. Our studies reveal how viral RNAs can use a combination of static and dynamic RNA structures to bind host machinery through highly noncanonical interactions and highlights the utility of cryo-EM for visualizing small conformationally dynamic structured RNAs.

## INTRODUCTION

RNA’s remarkable functional versatility derives from its unique ability to both encode genetic information and form complex three-dimensional structures (*1, 2*). RNA viruses universally exploit these features, using structured RNA elements to manipulate the host’s cellular machinery and regulate essential viral processes (*3–5*). Such RNA elements exist in viral clades as diverse as flaviviruses, lentiviruses, coronaviruses, alphaviruses, and picornaviruses (*3, 4, 6-12*), where they are often ‘multifunctional’ in that one RNA sequence performs more than one function. Despite their importance, our understanding of such RNAs is rudimentary, partly because many are conformationally dynamic and therefore difficult to characterize structurally using conventional methods. Crystallization of such elements is difficult due to their inherently dynamic structure, and nuclear magnetic resonance (NMR) is often not tractable for fully functional RNAs or RNA-protein complexes due to molecular weight limitations (*13, 14*). Thus, the structure-function rules that govern the molecular mechanism of multifunctional conformationally dynamic viral RNA elements, and their interactions with the cell’s machinery, are not well understood.

A powerful way that viruses use RNA structure is to mimic cellular transfer RNAs (tRNAs). For example, several viral internal ribosome entry sites (IRESs), including that of hepatitis C virus and viruses in the *Dicistroviridae* family, mimic parts of tRNA to contact tRNA binding sites on the ribosome and facilitate viral protein production (*3*). In addition, the 5’ untranslated region (UTR) of the HIV-1 RNA genome contains a tRNA-like element that binds lysyl-tRNA synthetase and favors the release of bound tRNA^Lys3^, the primer for HIV reverse transcription (*9*), and at least one DNA herpesvirus produces tRNA-like transcripts that may influence pathways linked to virulence (*15*). These and other examples show the importance of tRNA mimicry in diverse viruses, including some that cause human disease.

Important examples of tRNA mimicry and multifunctionality are the ‘tRNA-like structures’ (TLSs) at the 3’ ends of certain positive-strand RNA viral genomes (*16, 17*). TLSs were identified by their ability to drive aminoacylation of viral genomic 3’ ends by host aminoacyl-tRNA synthetases (aaRSs) and were later found to interact with other tRNA-specific enzymes including CCA-nucleotidyltransferase (CCA-NTase) and eukaryotic translation elongation factor 1A (eEF1A) (*16, 18*). Known TLSs are classified into three types based on their aaRS specificity: valylatable (TLS^Val^), histidylatable (TLS^His^), and tyrosylatable (TLS^Tyr^) (*16, 18*). Each type has a characteristic secondary structure that differs dramatically from tRNA, showing that putative tRNA mimicry can be achieved in diverse ways (Fig. S1) (*18*). Of the three classes, TLS^Tyr^ is the most different from tRNA in size and secondary structure and is the most difficult to reconcile with purported tRNA mimicry (*19*). The prototype TLS^Tyr^ is found at the 3’ end of each of the three genomic RNAs of the tripartite brome mosaic virus (BMV) genome (Fig. 1A). The BMV TLS is multifunctional, playing roles in translation, replication, and encapsidation of the BMV RNAs; some of these functions are linked to aminoacylation of the TLS, while other functions appear to be independent of aminoacylation status (*16, 19*). The BMV TLS is a thus powerful model system to explore features critical for function in diverse viral RNAs: tRNA mimicry, multifunctionality, host protein binding, and potentially conformational dynamics.

**Fig. 1.**
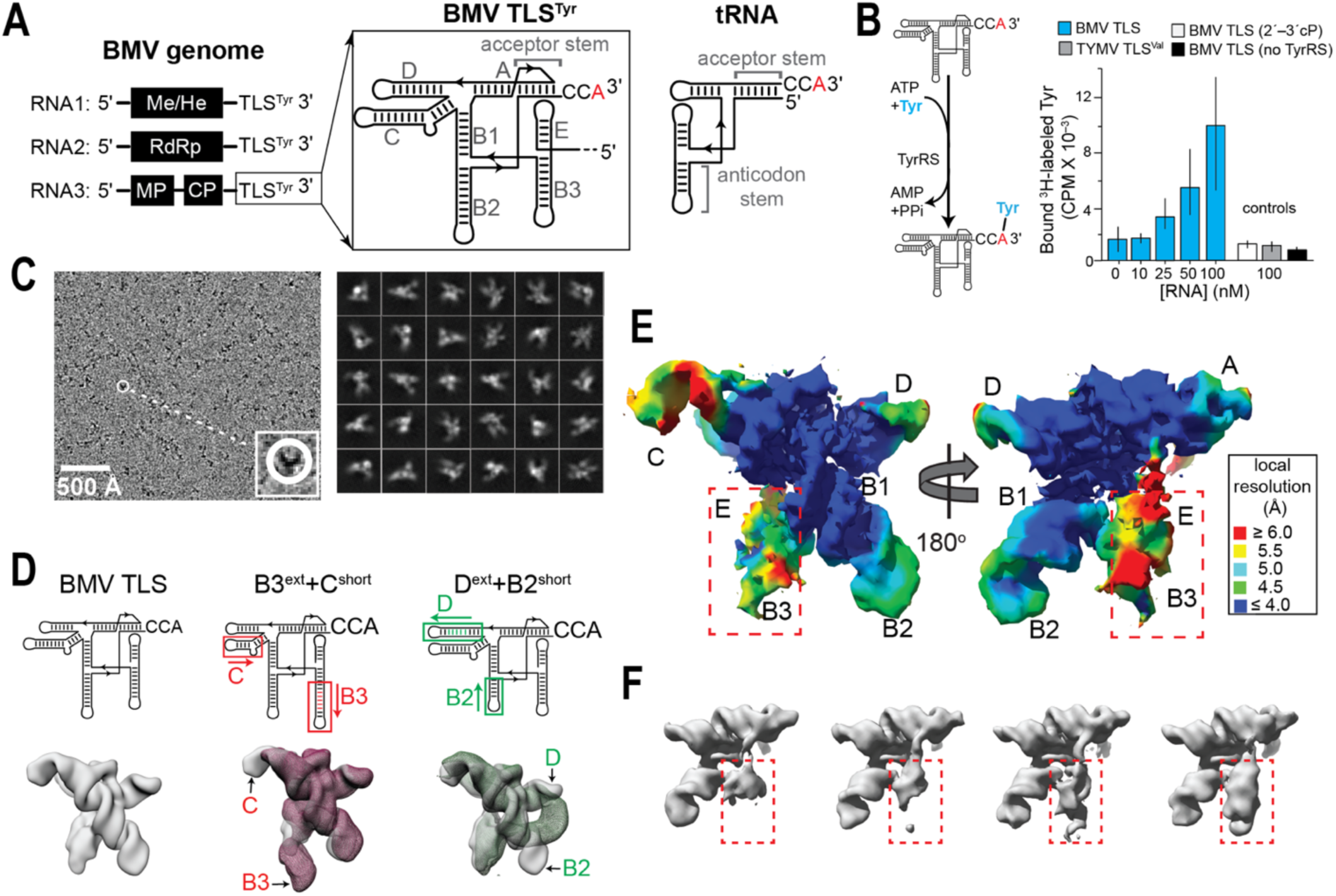
Functional and initial structural characterization of BMV TLS RNA. (A) Organization of the tripartite BMV genome with tyrosylatable TLS at the 3’ end of each RNA. Me/He: methyltransferase/helicase; RdRp: RNA-dependent RNA polymerase; MP: movement protein; CP: coat protein. The conserved secondary structure of the BMV TLS^Tyr^ is shown next to that of canonical tRNA. Structural domains and the terminal CCA are shown, with the red A designating the site of aminoacylation. (B) Left: Cartoon representation of chemical reaction catalyzed by TyrRS. Right: Tyrosylation of the BMV TLS RNA used for cryo-EM studies. Incorporation of radioactive tyrosine onto RNA was measured using an adapted published protocol (*47*). BMV TLS (2’–3’ cP) contains a terminal 2’–3’ cyclic phosphate and therefore it is not an efficient substrate of TyrRS. TYMV TLS^Val^ is a valylatable TLS from Turnip Yellow Mosaic Virus (TYMV). Aminoacylation reactions were performed for 30 minutes at 30 °C. The concentration of TyrRS was 100 nM. (C) Left: Representative micrograph of the BMV TLS^Tyr^ RNA with 30% of the field of view shown. Defocus range was –1 to –2.5 µm. Right: Classified projections from particles used to build initial 3D map of BMV TLS (Fig. S2). (D) Secondary structures and cryo-EM maps of BMV TLS, B3^ext^+C^short,^ and D^ext^+B2^short^. The maps of B3^ext^+C^short^ (mesh red) and D^ext^+B2^short^ (mesh green) are superimposed on BMV TLS (solid grey). Arrows point to differences in electron density that correspond to altered stems. (E) Refined map of BMV TLS generated using cryoSPARC with a resolution of 4.3 Å. The resolution was estimated using half maps and a gold standard FSC of 0.143 and was corrected for overfitting of high-resolution noise using noise substitution (*24*). Colors denote local resolution. Flexible domain with lower local resolution is inside red box. (F) Maps representing heterogeneity in the particles using 3D variability analysis in cryoSPARC (*25*). The variability among particles is mostly localized to the domain boxed in red.

Early studies identified the 3’-most 134 nucleotides (nts) of the genomic BMV RNAs as the minimal tyrosylatable TLS RNA (*19, 20*) but later studies demonstrated the importance of adjacent upstream sequence for efficient aminoacylation (*19*). The consensus sequence of 169 nts–i.e., the minimal tyrosylatable core plus flanking functionally important domains–forms a secondary structure with seven helical stems (compared to four for tRNA), including a pseudoknotted stem that serves as the aminoacyl acceptor arm analog (Fig. 1A; Fig. S1) (*18, 21*). Thus, BMV TLS is substantially larger and more structurally complex than tRNA and presents an example of highly divergent structure that accomplishes tRNA mimicry in a manner that is not readily apparent. It is not known whether domains within BMV TLS form a tRNA-like L-shape structure, and conflicting evidence points to either stem B2 or B3 as a potential anticodon stem analog (Fig. 1A) (*22*). Notwithstanding the lessons that a 3D structure of a TLS^Tyr^ could provide about viral RNA tRNA mimicry, multifunctionality, and co-opting of host cell machinery, the tertiary structure of BMV TLS has remained elusive and mysterious for decades.

To overcome this impasse, we used single-particle cryo-electron microscopy (cryo-EM) to explore the structure of BMV TLS, obtaining a map of the free unbound ∼55 kDa RNA to 4.3 Å. Unexpectedly, the elements involved in tRNA mimicry are not preorganized into a tRNA-like shape for binding TyrRS. Instead, by obtaining cryo-EM data of the BMV TLS-TyrRS complex (5.5 – 6.0 Å resolution), we show that binding TyrRS requires a large structural change, facilitated by inherent conformational dynamics, into a shape that bears little resemblance to tRNA. Further, our studies of the complex suggest that the TLS binds the TyrRS in more than one way. Despite this dynamic nature of the BMV TLS structure, the replication promoter is maintained in proximity to the replication initiation site in both unbound and bound states, suggesting how replication initiation may be facilitated by BMV TLS. Decades of functional and biochemical studies of the BMV TLS can now be reconciled and readily explained, including conflicting models about the identity of its tRNA-mimicking elements. More broadly, these results have implications for understanding tRNA mimicry, multifunctionality, RNA folding and dynamics, and RNA-protein interactions, and they also highlight the power of cryo-EM for visualizing the structure and inferring the mechanism of small dynamic functional structured RNAs and RNA-protein complexes once elusive to structural biology.

## RESULTS

### Cryo-EM reveals the global architecture of unbound BMV TLS

To obtain a functional folded BMV TLS RNA for cryo-EM studies, we *in vitro* transcribed and purified a previously characterized TLS sequence from BMV RNA 3 (*18, 21*) and confirmed its ability to be tyrosylated *in vitro* using recombinantly expressed TyrRS from model host *Phaseolus vulgaris* (Fig. 1B). Cryo-EM micrographs of this sample using a 200 kV microscope without a phase plate contained readily identifiable particles despite their small size (55 kDa) (Fig. 1C; Fig. S2). Using a standard data analysis pipeline that includes “junk” particle removal via 2D classification, *ab initio* 3D reconstruction using stochastic gradient descent optimization, 3D classification, and model refinement (*23*), we obtained an initial 7.0 Å map that displayed characteristics consistent with a folded RNA of the expected size (Fig. 1D; Fig. S2). The overall architecture of the map was robustly reproduced across *ab initio* reconstructions with different numbers of classes (Fig. S2).

A larger dataset from a 300 kV microscope equipped with an energy filter reproduced our initial results and increased the overall resolution to 4.3 Å. In this map, both minor and major grooves of A-form helices were clearly defined, and the connectivity of the phosphate backbone could be deduced (Fig. 1E; Fig. S3). In regions with relatively high local resolution, stacking and coplanarity of base pairs (bp) were resolved, and phosphate “bumps” were visible (Fig. S4). The central core of the map displayed the highest local resolution, but even peripheral region density was well-defined and displayed clear helical features (Fig. 1E).

Importantly, one helical domain stood out as it was less defined and had lower local resolution relative to other map regions (Fig. 1E, boxed), suggesting local flexibility within the structure. To examine potential conformational heterogeneity, we used 3D Variability Analysis (3DVA), a computational tool to dissect continuous conformational flexibility and/or multiple discrete conformations in a set of particles (Fig. 1F) (*25*). A series of 3D volumes generated by this analysis represents the variability among particles within the dataset. This analysis was consistent with this one helical domain occupying multiple conformational states, while the rest of BMV TLS RNA is relatively more static (Fig. 1F).

### Engineered RNAs provide information for unambiguous structural modelling

Aside from a low-resolution small angle X-ray scattering (SAXS) envelope and a computational model based on chemical probing and functional data (*18, 26*), there was no prior information on the 3D structure of BMV TLS. To assign RNA helices to regions of the cryo-EM map, we used a strategy based on the well-established concept of RNA structural modularity. Specifically, we designed and *in vitro* transcribed BMV TLS RNAs with certain helical stems extended or truncated by several base pairs, with the rationale that local differences between the wild type and modified RNA constructs would allow identification of the extended/truncated helices and assignment of the density. Analogous experiments have been used in previous studies to validate structural models or to measure the angles between helices emerging from an RNA catalytic core (*27, 28*).

A mid-resolution map of an RNA with stem B3 extended and stem C shortened displayed local differences compared to wild type that were consistent with the extension of one helix and truncation of another, thus conclusively identifying B3 and C (Fig. 1D; B3^ext^+C^short^). We repeated this analysis with an RNA with stem D extended and stem B2 shortened (Fig. 1D; D^ext^+B2^short^). An unexpected new tertiary interaction apparently formed between the modified helices, but this did not affect the global architecture of the RNA, and local differences between the maps identified stems B2 and D. This strategy, in combination with the previously determined secondary structure (*26*) (Fig. 2A), allowed us to unambiguously identify the positions of all helices with no prior assumptions. This method of obtaining RNA helical assignments in cryo-EM maps is generally applicable and can also validate structural models from automated computational tools (*29*), as discussed below and done previously (*27*).

**Fig. 2.**
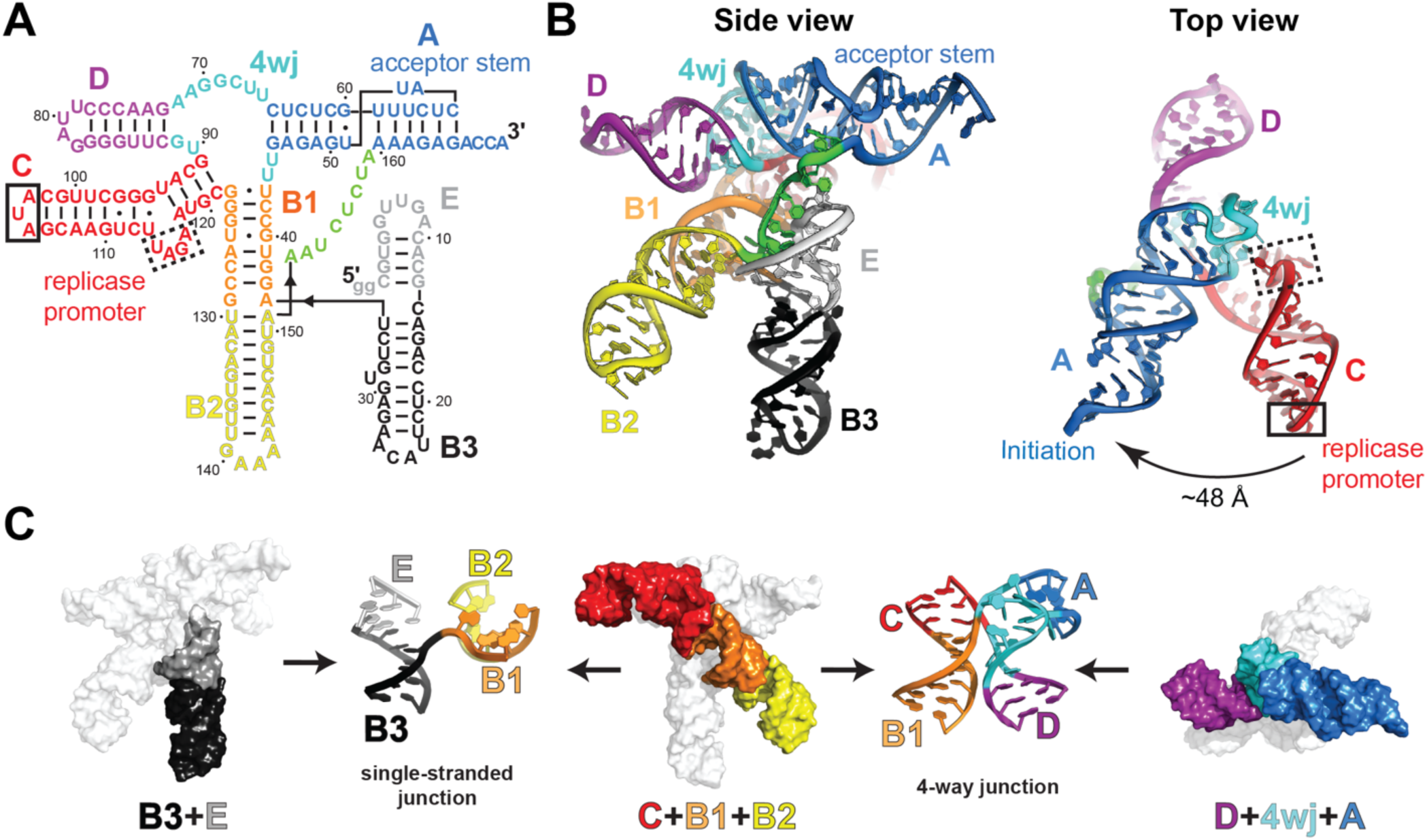
Structure of free BMV TLS^Tyr^ RNA. (A) Secondary structure of the BMV TLS^Tyr^ RNA construct used for cryo-EM is labeled and colored by domain, with the locations of the replicase promoter and the acceptor stem labeled. The AUA apical triloop (solid box) and UAGA internal loop (dashed box) within stem C are critical for replication. (B) Structural model of BMV TLS viewed from side and top views. Colors match those in the secondary structure. In top view, sequences that are critical for replication are boxed as in (A). (C) The helices comprising three domains of BMV TLS^Tyr^ formed by helical stacks are highlighted with their corresponding colors: B3+E (left, black and grey), C+B1+B2 (middle, red, orange and yellow), and D+4wj+A (right, purple, cyan and blue). The junctions connecting the different domains are shown between them. Colors match those in (A) and (B).

### Cryo-EM yields a complete structural model of BMV TLS

To build a structural model that is consistent with the cryo-EM data and the previously determined secondary structure of BMV TLS, we evaluated a decades-old computational 3D model (Fig. S5) (*26*). The computational model correctly predicted five helical stems emanating from a central core but did not fit well into the experimental map. We built a new structure by docking individual domains into their corresponding density, performing molecular dynamics flexible fitting and real space refinements with Phenix, and correcting RNA geometry using tools based on Rosetta modelling (Fig. 2B; Fig. S6) (*30–33*). The entire structure could be built and refined within the map without substantial steric clashes or breaks in the chain (Table S1). Atomic models of the extended/truncated constructs (Fig. 1D) support this structural model of BMV TLS (Fig. S7).

As a complementary method of structural modelling, we used auto-DRRAFTER, a fully automated computational tool that generates multiple unbiased models from a user-provided secondary structure and low-to-moderate resolution cryo-EM maps (*29*). The models generated by auto-DRRAFTER agreed very well with our original structure (RMSD < 3.2Å) and with the helical assignment information obtained from the extensions/truncations (Fig. 1D; Fig. S6B).

Globally, the BMV TLS structure contains three sets of coaxially stacked extended helical domains (Fig. 2C). One domain comprises helices C, B1, and B2, which form a pseudo-continuous helix that spans the length of the structure, with the apical loops of B2 and C pointing in opposite directions (Fig. 2C, center). The second comprises helices D and A connected through a central 4-way junction (4wj; Fig. 2C, right). Helix A contains the 3’ CCA and serves as both the acceptor stem for tyrosylation and the replication initiation site (*16*). The third corresponds to the conformationally dynamic domain mentioned above (Fig. 1F) and contains helices B3 and E (hereafter referred to as B3+E) linked to the core of the structure by a single strand of unpaired RNA (Fig. 2C, left). Thus, while atomic-level details are ambiguous at the overall map resolution, the cryo-EM structure of BMV TLS reveals functionally important features.

### Identification of elements that drive aminoacylation

The ability of the BMV TLS to be aminoacylated has led to the hypothesis that the “rules” for BMV TLS tyrosylation by TyrRS largely match those for tRNA^Tyr^ (*26, 34*). Specifically, because tRNA^Tyr^ recognition by the aaRS requires interactions with its acceptor stem and anticodon loop (with the acceptor stem being more important), analogs of these domains were expected in the BMV TLS structure (*16, 26, 35, 36*). Pseudoknotted helix A was clearly identified as the acceptor stem analog in previous studies, but the identity of the putative anticodon stem analog has remained a mystery (*16, 37*), with conflicting evidence for stems B3 or B2 (*22, 26, 34, 37*). Our 3D structure shows that while B2 is oriented away from the acceptor stem, the B3+E domain is positioned such that it is more likely to interact with TyrRS (Fig. 2B). Furthermore, a consensus sequence and secondary structural model based on conservation and covariation analysis of 512 unique viral BMV-like TLS^Tyr^ sequences reveals that certain nucleotide identities in B3’s apical loop and the length of the B3+E domain are highly conserved (Fig. 3A & B), suggesting a selective pressure constraining these characteristics. In contrast, stem B2 varies in length with no compensatory changes in the length of B1 (Fig. 3B), and there is no strong conservation in the apical loop of B2 that would suggest that it acts as the anticodon loop (Fig. 3A). The secondary structure model derived from the covariation analysis was validated with chemical probing of three representative TLS variants (Fig. S8). These observations strongly point to B3 as the best candidate for the putative anticodon stem analog.

**Fig. 3.**
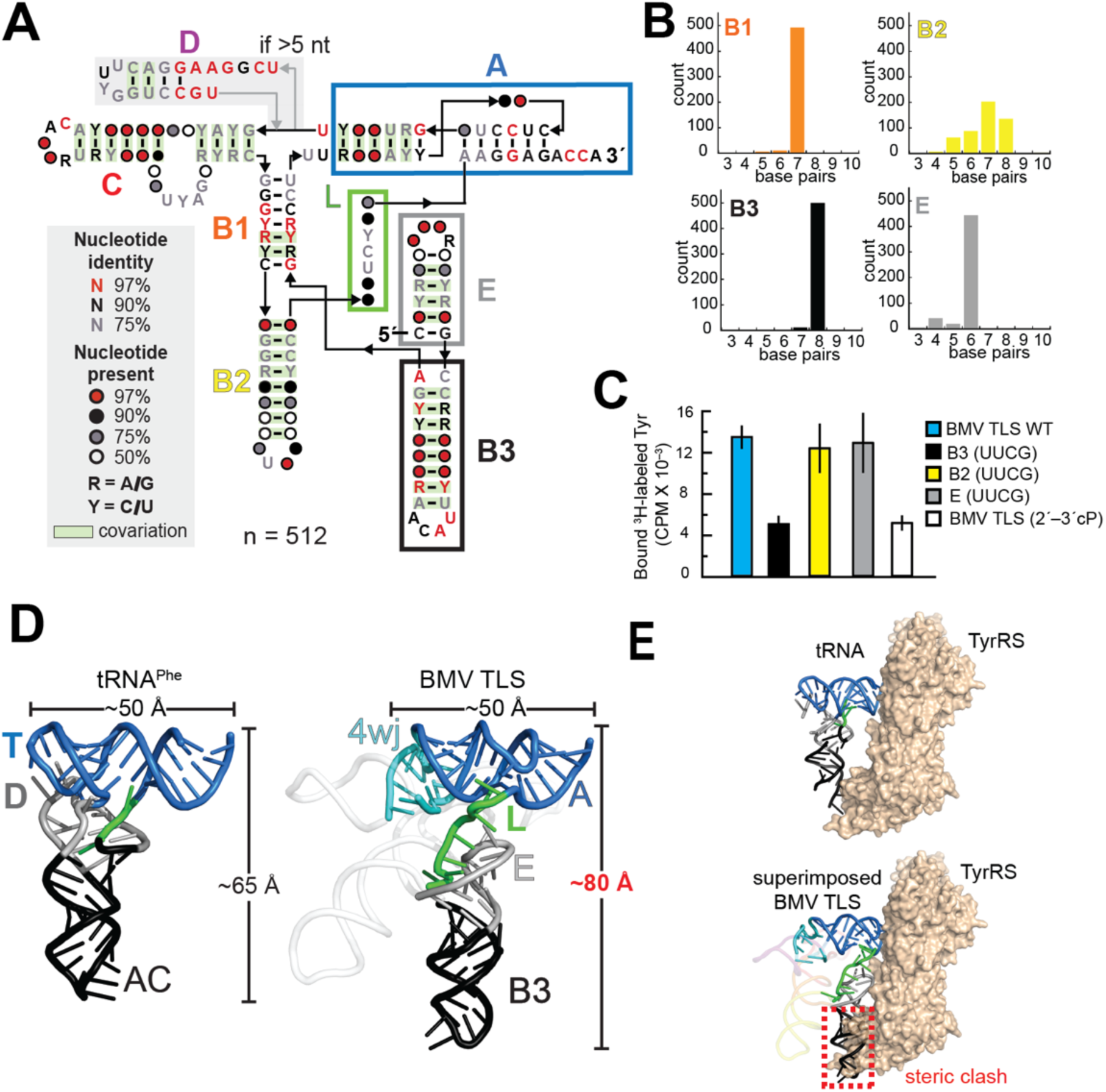
Structural features of BMV TLS^Tyr^ important for replication or tRNA mimicry. (A) Consensus covariation and sequence model of TLS^Tyr^ variants related to BMV TLS. Helices are labeled according to BMV TLS. The model began with a preliminary alignment of eight TLS^Tyr^ sequences from the Rfam database (*49, 50*), followed by homology searches that resulted in 512 unique sequences then used to build a consensus model. The secondary structural model is supported by SHAPE chemical probing of three representative variants from different viruses (Fig. S9). (B) Distribution of lengths of four helices according to the structural alignment of 512 sequences. The high conservation of the lengths of E and B3 supports the role of these helices as the anticodon stem analog in BMV TLS. (C) Tyrosylation of mutant BMV TLS described in the main text. BMV TLS (2’–3’ cP) contains a terminal 2’–3’ cyclic phosphate and therefore it is not an efficient substrate of TyrRS. Aminoacylation reactions were performed for 5 minutes at 30 °C. The concentrations of RNA and TyrRS were 120 and 100 nM, respectively. (D) Comparison of the structure of unbound tRNA^Phe^ (left) and tRNA mimicking portions of the BMV TLS^Tyr^ (right). The structurally important loops T and D are labeled in tRNA^Phe^, as is the anticodon (AC) loop. Analogous structural features are colored as per Fig. 2 and various molecular dimensions are shown. (E) (top) Published crystal structure of yeast tRNA^Tyr^ bound to yeast TyrRS (*36*). (bottom) The cryo-EM-derived BMV TLS structure overlaid on the bound tRNA^Tyr^ by superimposing the acceptor arms of each (Fig. S11). The location of a steric clash between B3 (anticodon stem analog) and TyrRS is boxed in red.

To further investigate the role of B3 in tRNA mimicry by BMV TLS, we performed *in vitro* tyrosylation studies of BMV TLS variants in which the terminal loops of B2, B3, or E were mutated to UUCG, a sequence used to cap helices due to its well-determined structure and high thermodynamic stability (*38*). In contrast to large sequence deletions used in previous studies (*22*), the UUCG mutations were not expected to affect overall BMV TLS structure. Indeed, electrophoresis of folded BMV TLS mutants under native conditions showed unchanged migration relative to wild type (Fig. S9). While mutations to B3 had a clear detrimental effect on aminoacylation of the TLS (Fig. 3C), mutations to B2 or E did not have a substantial effect, suggesting B3 is analogous to the tRNA anticodon. Although the B3 apical loop contains a conserved ‘UACA’ sequence rather than a canonical tyrosine anticodon (i.e. GUA in plants), the major identity elements of tRNA^Tyr^ lay within its acceptor arm, and interactions between TyrRS and the anticodon play a less important role (*35*); this appears also to be true with the BMV TLS.

### Free BMV TLS RNA does not contain a classic L-shape tRNA mimic

To productively interact with the synthetase, it was expected that BMV TLS’ acceptor and anticodon stem analogs (stems A and B3, respectively) would comprise a classic L-shaped fold of similar size and shape to tRNA (Fig. 3D, left). Surprisingly, stem A and domain B3+E of BMV TLS are only loosely associated, with no interactions analogous to those between the T and D loops of tRNA (Fig. 3D, right). Thus, the overall dimensions of the tRNA-mimicking portions of the BMV TLS do not match tRNA (Fig. 3D). The implications were apparent when we used the published crystal structure of a yeast tRNA^Tyr^-TyrRS complex (*36*) to model BMV TLS bound to TyrRS (Fig. 3E). Superposition of the structure of BMV TLS on tRNA^Tyr^ bound to the TyrRS homodimer, based on alignment of their acceptor stems (Fig. S10), revealed substantial steric clashes between TyrRS and the B3+E domain of BMV TLS (Fig. 3E; Fig. S10B). This suggests that BMV TLS may require conformational changes to bind TyrRS, and/or binds TyrRS with a geometry that differs from canonical tRNA, and/or binds to a different binding site. Indeed, the fact that B3+E is conformationally dynamic made it plausible that interaction with TyrRS requires a rearrangement of this domain.

### BMV TLS undergoes large conformational changes to bind TyrRS

To examine if the structure of the BMV TLS must change to productively bind TyrRS, we applied cryo-EM to the BMV TLS-TyrRS complex reconstituted from pure components (Fig. 4; Fig. S11 & S12). Initial studies with a 200 kV microscope (Fig. 4A) clearly showed two copies of BMV TLS RNA (Fig. 4B, red arrows) bound to opposite sides of the TyrRS homodimer (Fig. 4B, cyan arrow), with each BMV TLS RNA making two contacts with the enzyme, presumably using the acceptor and anticodon stems. This resembles the overall configuration of the tRNA^Tyr^-TyRS complex (Fig. 4C), but further inspection suggested distinct binding modes. Specifically, there is significantly more space between the TLS and the surface of the enzyme compared to tRNA^Tyr^-TyrRS, suggesting that the angles between the acceptor and anticodon stems differ between the bound TLS and tRNA^Tyr^. Consistent with our interpretation of the density, cryo-EM data of free TyrRS (Fig. S13) matched the density observed in the center of the BMV TLS-TyrRS complex (Fig. 4B, cyan box).

**Fig. 4.**
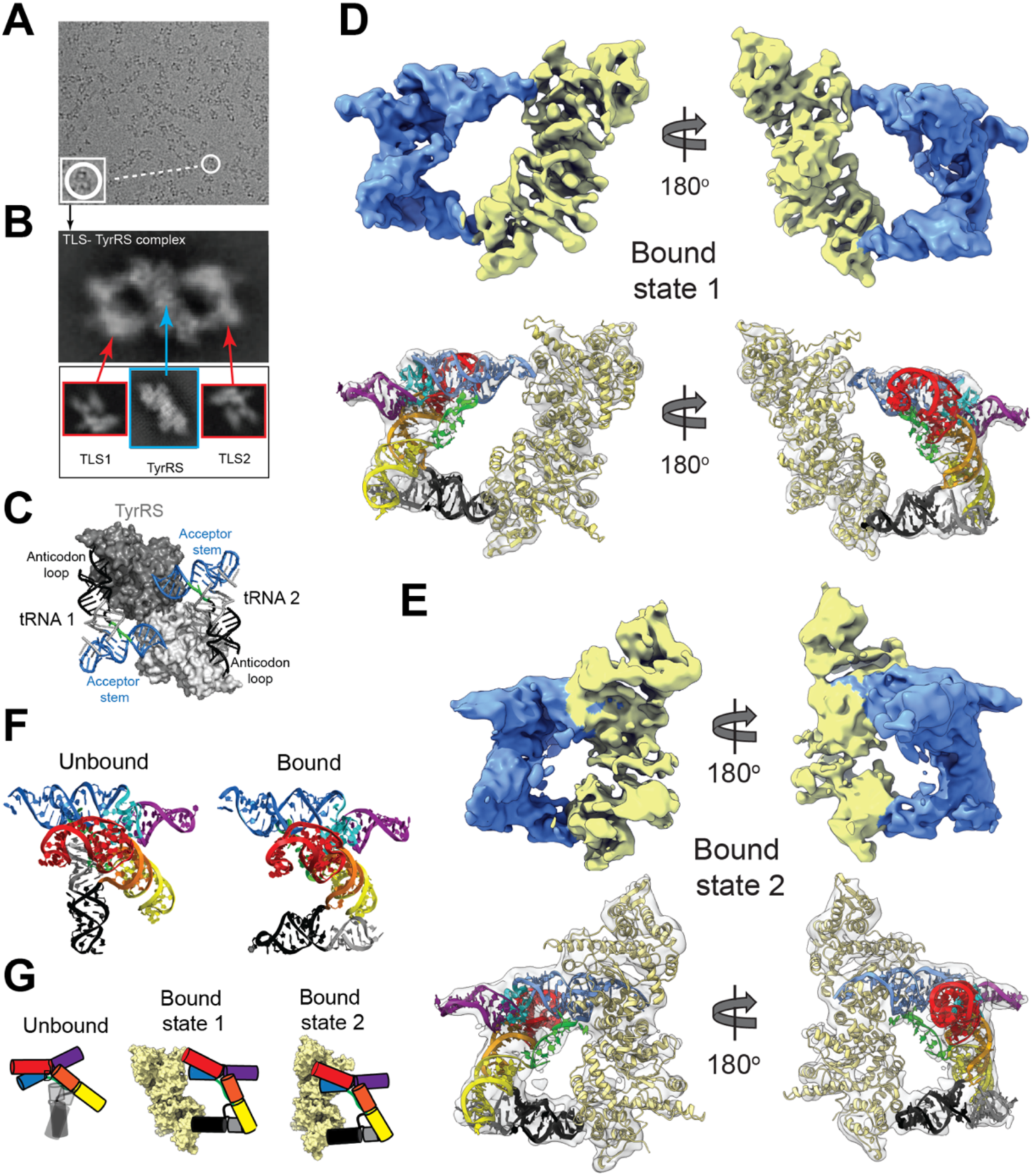
Cryo-EM studies of the BMV TLS-TyrRS complex. (A) Representative micrograph of the BMV TLS-TyrRS complex with a 200 kV electron microscope. The complex was reconstituted from purified recombinant TyrRS from *Phaseolus vulgaris* (common bean), a model host of BMV, and *in vitro* transcribed BMV TLS RNA. (B) Representative 2D class of BMV TLS-TyrRS complex showing two BMV TLS RNA molecules bound to TyrRS. For comparison and interpretation of the density of the complex, densities of unbound TyrRS (cyan box; Fig. S13) and unbound BMV TLS RNAs (red boxes; Fig. 1C) are shown on the bottom. (C) Crystal structure of the yeast tRNA-TyrRS complex, with tRNA elements labeled (PDB ID: 2dlc). (D) A 5.5 Å cryo-EM map of the BMV TLS-TyrRS complex in ‘bound state 1’ (top) and an atomic model fitted to the density (bottom). The structure of *Phaseolus vulgaris* TyrRS was built using homology modelling. (E) A 6.0 Å cryo-EM map of the BMV TLS-TyrRS complex in ‘bound state 2’ (top) and an atomic model fitted to the density (bottom). (F) Comparison of the structure of BMV TLS in isolation vs. bound to TyrRS. (G) Comparison of BMV TLS in unbound state and two bound states. In the unbound state the RNA exists as a dynamic conformational ensemble in which the anticodon stem analog samples multiple conformations. Binding to TyrRS stabilizes the anticodon stem in a specific orientation that is roughly parallel to that of the acceptor arm in both bound conformations. The two bound states differ in the relative positions of BMV TLS RNA and TyrRS. Colors are as in Fig. 2.

To learn more about the binding behavior of BMV TLS, we collected a larger dataset with a 300 kV microscope at different tilt angles to reduce the effect of preferred particle orientations (Fig. S11). Although two bound BMV TLS RNAs are observed in many of the 2D classes (Fig. 4B; Fig. S11), one RNA was consistently better defined than the other in the 3D reconstructions. This suggests that the complex is conformationally dynamic and/or there is compositional/conformational heterogeneity. Consistent with the latter, and as discussed below, the data are consistent with at least two distinct bound conformational states, ‘bound state 1’ and ‘bound state 2’ (Fig. 4D-E; Fig. S11). Interestingly, a previous study reported that although the TyrRS homodimer has two tRNA^Tyr^ binding sites and crystallizes with two copies of tRNA^Tyr^, only one tRNA^Tyr^ is aminoacylated at a time in solution (*39*). We speculate that this apparent paradox may be reflected in the asymmetrical binding behavior of the two BMV TLS RNAs, but further studies are required.

Focusing on a single copy of BMV TLS RNA bound to TyrRS, we obtained interpretable cryo-EM 3D maps that clearly show the global conformation of the TLS when bound to TyrRS (Fig. 4D & E, top panels; Fig. S12), with resolutions of 5.5 and 6.0 Å for bound states 1 and 2, respectively (Fig. S12). The maps agreed with the molecular dimensions of TyrRS and BMV TLS RNA and displayed clear secondary structural features, allowing us to fit atomic models of BMV TLS RNA and TyrRS (Fig. 4D & E, bottom panels).

The overall conformation of BMV TLS is essentially the same in both bound states (Fig. S14A). Importantly, when compared to the conformation of the unbound TLS, the structures of the complex show that BMV TLS RNA undergoes a large conformational change to bind TyrRS (Fig. 4F). Specifically, the structures of the unbound and bound BMV TLS RNA were essentially superimposable except for domain B3+E (Fig. S14B). Thus, in the unbound state, B3+E occupies a conformational ensemble, mostly occupying a position at roughly a right angle to the acceptor stem but not properly positioned to interact productively with the enzyme (Fig. 3E). However, in the bound state B3+E has rotated ∼90° from its average unbound position to lay roughly parallel to the acceptor stem analog (stem A), avoiding any steric clash with the enzyme and placing the B3 apical loop on the surface of TyrRS (Fig. 4D-F). The bound BMV TLS geometry is very different from tRNA^Tyr^ in that while BMV TLS also makes two discrete contacts with TyrRS, it has little resemblance to the global L-shaped structure of tRNA^Tyr^. This bound conformation was not observed in our analysis of the free RNA, suggesting it is unstable and rarely adopted in the absence of interactions with TyrRS. Thus, the BMV TLS must undergo a dramatic programmed conformational change to bind the synthetase, in contrast to preorganized tRNAs and TYMV TLS^Val^; the latter nearly perfectly mimics tRNA (*40*).

### BMV TLS binds TyrRS in at least two distinct states

As mentioned in the previous section, we observed the BMV TLS-TyrRS complex in two distinct states that may relate to the process of aminoacylation. The overall conformations of the RNA and the enzyme appear to be the same in both states (Fig. S14A & C), but their relative positions are different. Specifically, the anticodon loop analog is in approximately the same position in both states, but the position of the acceptor stem changes (Fig. 4D & E). In bound state 1 the acceptor stem makes limited contacts with TyrRS and the terminal CCA is well outside the aminoacylation active site (Fig. 4D). In bound state 2, the RNA and the enzyme are closer, the acceptor stem makes deeper contacts with TyrRS, and the terminal CCA is positioned in the active site. Because the position of the acceptor stem in bound state 2 more closely resembles that of tRNA^Tyr^ bound to TyrRS, this state is more likely to reflect the bound conformation during aminoacylation. However, we cannot make conclusions about the order of events leading to aminoacylation, for example, whether bound state 1 is an ‘on path’ intermediate or an alternate non-productive state. Additionally, as the conformation of the second bound RNA was not resolved, we do not know if all possible combinations of bound states 1 and 2 are present in the sample, or if there are preferred combinations. Intriguingly, while this behavior may be unique to BMV TLS, it is possible that similar multiple bound states are occupied during aminoacylation of authentic tRNA^Tyr^ by TyrRS but only a single state was observed by crystallography. Cryo-EM studies of tRNA^Tyr^-TyrRS and further functional and structural characterization of TLS^Tyr^-TyrRS complexes may provide additional insights.

### The replicase promoter is pre-positioned in proximity to the initiation site

A critical function of BMV TLS is to recruit the replication machinery to the proper initiation site (*19, 41, 42*). Although replication initiates at the 3’ end in stem A, the replicase promoter is within stem C in the apical AUA triloop (Fig. 2A & B, solid box) and a UAGA 4-nt bulge (Fig. 2A & B, dashed box) (*41–43*). The structure revealed that helix C is prepositioned adjacent and nearly parallel to helix A (Fig. 2B, right). The resulting distance between the AUA triloop promoter and the replication initiation site is ∼48 Å. As the replication complex consists of multiple viral and host proteins ranging in size from 43 to 110 kDa (BMV encoded replication proteins P1 and P2 are 110 and 95 kDa, respectively) (*44*), the replication complex is likely large enough to span this distance. Further, the proximity of the UAGA 4-nt bulge (Fig. 2B, dashed box) to the 4wj likely creates tertiary interactions that stabilize the orientation and positioning of stem C relative to helix A, and this may be part of that sequence’s role (Fig. 2B, right). Overall, the structure suggests that the relative position of helix C and helix A is preformed to reduce the conformational search of the bound replicase for its substrate. However, we cannot exclude the possibility that binding of the replicase, as with TyrRS, is associated with RNA conformational changes.

## DISCUSSION

The BMV TLS has served as a model system for understanding RNA structure-function relationships for decades, but its three-dimensional structure remained elusive. Using cryo-EM we have elucidated the tertiary conformation of BMV TLS and revealed not only structural features, but previously unknown conformational dynamics that may explain why crystallographic efforts failed. The structure of the BMV TLS in the unbound and TyrRS-bound states reveals a novel strategy for co-opting the cell’s machinery using an RNA structure that contains a combination of conformationally dynamic and relatively static elements. Remarkably, the BMV TLS achieves aminoacylation without directly mimicking the overall 3D structure of a tRNA in either the unbound or bound forms; the decades-old term ‘tRNA-like structure’ may perhaps be a misnomer when applied to this RNA. Rather, the CCA at the 3’ end and an anticodon loop analog in B3 are positioned within an architecture that has little resemblance to canonical tRNA, but which spatially arranges them to interact with the TyrRS. This surprising mode of binding invites speculation that other RNAs with secondary structures that deviate dramatically from tRNAs may be capable of binding aaRSs in non-canonical ways; these would be difficult to identify based on sequence or secondary structure.

The potential function and/or consequences of BMV TLS’ conformational changes and a global fold that differs from tRNA are not clear. It could be that this conformational change serves as a signal to initiate synthetase binding or enables the binding of other proteins to the remodeled structure. We note that in the unbound form, the apical loop of stem E is occluded, but it is exposed in the bound form and in proximity to stem B2; the functional implications of this, if any, are unknown. The programmed conformationally dynamic nature of the RNA structure might also help confer multifunctionality. As described above, the BMV TLS interacts with various other host and viral proteins to function in replication, recombination, and encapsidation of the viral RNAs (*41*). It is possible that BMV TLS undergoes conformational changes to accommodate interactions with the distinct machineries required for each of these functions and that a static preformed structure would limit this. We note, however, that the replication promoter and initiation sites are proximally pre-positioned; hence the structure is a combination of preformed and dynamic features. Other multifunctional RNAs likely utilize similar combinations of dynamic and relatively static structural features to organize their different roles.

RNAs exist as conformational ensembles, and dynamics have emerged as a critical aspect of RNA function (*1, 13, 45*). Recent studies have already explored cryo-EM as a powerful tool for rapidly solving the structure of small RNA-only structures (*29, 46*) and our studies further highlight the potential of cryo-EM for dissecting not only structures, but dynamic processes involving functional structured RNAs and RNA-protein complexes. Unlike crystallographic studies, in which conformational dynamics must largely be inferred or suppressed, cryo-EM offers the advantage of direct detection of inherent motions. Here, we highlighted a tool that can aid in this task: the use of extensions and/or truncations of modular helical domains in combination with robust secondary structures to rapidly assign specific secondary structural elements to the electron density, provide constraints for structural modelling, and/or validate automated modelling programs. Cryo-EM, in combination with emerging computational tools, will greatly facilitate the study of diverse dynamic regulatory RNAs and RNA-protein complexes containing viral RNAs, riboswitches, ribozymes, aptamers, and many others.

## Acknowledgments

Eduardo Romero Camacho and Peter Van Blerkom (Univ. of Colorado Anschutz Medical Campus Cryo-EM Facility), and Theo Humphreys (Pacific Northwest Center for Cryo-EM) assisted with microscope operation. Erik Hartwick helped with the aminoacylation assays. The authors thank current and former Kieft Lab members for thoughtful discussions and technical assistance, and Anna-Lena Steckelberg, Benjamin Akiyama, Quentin Vicens, and David Constantino for critical reading of the manuscript.

## Funding

This work was supported by NIH grants R35GM118070 (JSK) and F32GM139385 (SLB). SLB is a Howard Hughes Medical Institute Hanna Gray Fellow. MES is a Jane Coffin Childs Postdoctoral Fellow. A portion of this research was supported by NIH grant U24GM129547 and performed at the PNCC at OHSU and accessed through EMSL (grid.436923.9), a DOE Office of Science User Facility sponsored by the Office of Biological and Environmental Research.

## Author contributions

SLB conceptualized the study, performed cryo-EM and aminoacylation experiments, analyzed data, performed structural modelling, and wrote the manuscript in collaboration with JSK. MES performed bioinformatic analysis and chemical probing and edited the manuscript. AM purified proteins and edited the manuscript. JSK conceptualized the study, performed structural modelling, acquired funding, and wrote the manuscript in collaboration with SLB.

## Competing interests

The authors declare no competing interests.

## Data and materials availability

All data are available in the manuscript and supplementary materials. Maps and atomic coordinates will be deposited in public databases before publication.

## Supplementary Materials for

### Other Supplementary Materials for this manuscript include the following

Supplementary_file_1.sto: Alignment of TLS^tyr^ variants used in consensus model.

Supplementary_file_2.xlsx: Organisms and accession numbers used in consensus model.

## MATERIALS AND METHODS

### *In vitro* transcription, purification, and folding of RNA

Templates for *in vitro* transcription were generated by PCR amplification of sequence-verified double-stranded DNA fragments (gBlocks; Integrated DNA Technologies). Sequences for all constructs are provided in Table S4. To reduce the amount of N+1 products generated by *in vitro* transcription and ensure correct termination at CCA_OH_, which is necessary for aminoacylation, PCR amplification was performed using a reverse primer with a 2′OMe modification at the second 5′ nucleotide, as described previously (*61*).

The gBlocks were synthesized with a ‘GG’ added at the 5’ end of the sequences of interest to facilitate transcription by T7 RNA polymerase. The transcription reactions contained 8 mM of each NTP (ATP, UTP, CTP, and GTP), 60 mM MgCl_2_, 30 mM Tris, pH 8.0, 10 mM DTT, 0.1% spermidine, 0.1% Triton X-100, 0.24 U/µl RNasin RNase inhibitor (Promega), and ∼ 0.14 mg/ml T7 RNA polymerase (in-house prepared). The transcription reactions were incubated for 3 hrs at 37°C. For constructs with 5′ and/or 3′ self-excising ribozymes (Table S4), the concentration of MgCl_2_ was increased to 120 mM after transcription and the constructs were incubated for an additional 15 minutes at 60°C. The transcribed RNA constructs were ethanol precipitated, purified by denaturing gel electrophoresis (8% polyacrylamide), cleaned using AMPure XP magnetic beads (Beckman Coulter), and buffer exchanged into RNase free water using 10 kD cutoff Amicon centrifugal filters (Millipore).

As one of the negative controls for BMV TLS aminoacylation, we produced BMV TLS (2’–3’ cP), which is identical to BMV TLS except for a terminal 2′-3′ cyclic phosphate that prevents efficient aminoacylation. To introduce the terminal 2′-3′ cyclic phosphate, a 69 nt hepatitis delta virus (HDV) self-excising ribozyme was introduced 3′ of the wild type BMV TLS sequence. BMV TLS (2’–3’ cP) was transcribed, cleaved, and purified using the protocol described above. The product of the ribozyme cleavage results in the production of the terminal 2′-3′ cyclic phosphate.

To fold the RNAs into their native conformation, the RNAs were resuspended at the desired concentration into a buffer solution containing 50 mM MOPS pH 7.0, 30 mM Na^+^, and 10 mM MgCl_2_, incubated at 80-90°C for 1 minute and cooled down to 23°C for 15 minutes.

### Preparation of TyrRS for aminoacylation assays and cryo-EM studies

To test the aminoacylation capacity of our BMV TLS RNA constructs and for cryo-EM studies of TyrRS and of the BMV TLS-TyrRS complex, we used TyrRS from a model host of BMV, *Phaseolus vulgaris* (common bean), as previous studies have shown that this efficiently tyrosylates BMV RNA *in vitro* (*62*). We obtained the sequence of this TyrRS from the UniProt database (gene: PHAVU_002G027700g) and optimized it for expression in *E. coli* using the online Integrated DNA Technologies (IDT) Codon Optimization Tool. A gBlock encoding the TyrRS sequence, cleavage sequences for restriction enzymes NdeI and BamHI, and a 6-His-tag was cloned into a pET15b vector using standard molecular biology techniques for ligation with T4 DNA ligase. The protein was expressed in BL21 (DE3) cells in Luria broth (LB) containing ampicillin at 37°C until OD600 reached 0.6 and then protein production was induced with the addition of 0.5 mM IPTG for 4 hours. The cells were harvested by centrifugation at 5488 x g for 12 minutes at 6°C. The pellets were resuspended in lysis buffer (20 mM Tris-HCl pH 7.0, 500 mM NaCl, 2 mM β-mercaptoethanol, 10% glycerol, and 1 EDTA-free protease inhibitor tab (Roche)) and sonicated for a total of 2 minutes (20 sec on and 40 sec off intervals) at 75% amplitude. The lysate was centrifuged at 30,000 x g for 30 min at 4°C before being loaded into a gravity flow column (Bio-Rad) and purified by Ni-NTA resin (UBP Bio) followed by size exclusion using FPLC and a Sepax 300 SEC column. The TyrRS protein was stored at a concentration of 10 µM (based on MW of dimer) in buffer containing 20 mM Tris-HCl pH 7.5, 500 mM NaCl, 2 mM β-mercaptoethanol, and 10% glycerol.

### Aminoacylation assays

#### Aminoacylation of sample used for cryo-EM studies

Tyrosylation of BMV TLS and control RNAs was assayed using a protocol adapted from a previous study of TYMV TLS^Val^ (*61*). Briefly, the RNAs were folded by heating and cooling in a buffer containing 10 mM MgCl_2_, 30 mM Na^+^, and 50 mM MOPS pH 7.0, as described above. The RNAs were diluted to 10 X the final reaction concentrations (0 – 100 nM; Fig. 1B). Prior to the aminoacylation experiment, the TyrRS was exchanged into a buffer containing 50 mM Tris pH 8.0, 5 mM MgCl_2_, 1 mM TCEP, and 5% glycerol using an Amicon 30 kD cutoff centrifugal filter and diluted to 1 µM (based on the MW of the dimer). The reaction buffer consisted of 2 mM ATP, 30 mM KCl, 5 mM MgCl_2_, 5 mM DTT, and 30 mM HEPES-KOH pH 8.0. The total volume of each aminoacylation reaction was 20 µl. Reaction buffer, TyrRS solution (final concentration of TyrRS was 100 nM), and folded RNA solution (final concentration of RNA was 0 – 100 nM) were mixed and incubated on ice. The aminoacylation reaction was initiated by the addition of 1 µl of ^3^H-labeled L-tyrosine (40 – 60 Ci/mmol) and followed by incubation at 30°C for 30 min. The reactions were then diluted by adding 200 µl of wash buffer (20 mM Bis-tris pH 6.5, 10 mM NaCl, 1 mM MgCl_2_, and trace amounts of bromophenol blue dye) and immediately deposited in a well of a vacuum filter blotting apparatus. The reaction solution was filtered through a layer of Tuffryn membrane (PALL Life Sciences), Hybond positively charged membrane (GE Healthcare), and gel dryer filter paper (Bio-Rad). The negatively charged RNA binds to the Hybond membrane. Immediately after the diluted reaction solution ran through the filter membranes, 600 µl of wash buffer was added to the well and allowed to flow through the membranes. The Hybond membrane was allowed to dry and the locations marked by the bromophenol blue from the wash buffer were cut out. A scintillation counter (Perkin-Elmer Tri-Carb 2910 TR) was used to measure the radioactivity of the Hybond membrane pieces.

#### Comparison of BMV TLS UUCG mutants

Wild type BMV TLS and mutants with UUCG at the apical loops of B3, B2, or E (Table S4) were transcribed and purified as described above. The RNAs were folded in 10 mM MgCl_2_, 30 mM Na^+^, and 50 mM MOPS pH 7.0 at a concentration of 1.2 µM. Three microliters of TyrRS stock solution ([TyrRS dimer] = 10 µM in 20 mM Tris-HCl pH 7.5, 500 mM NaCl, 2 mM β-mercaptoethanol, and 10% glycerol) was added to 27 µl of buffer containing 50 mM Tris pH 8.0, 5 mM MgCl_2_, 1 mM TCEP, and 5% glycerol. A 10X reaction buffer consisted of 20 mM ATP, 300 mM KCl, 50 mM MgCl_2_, 50 mM DTT, and 300 mM HEPES-KOH pH 8.0. The total volume of each aminoacylation reaction was 20 µl. For each reaction we added 13 µl of water, 2 µl of 10X reaction buffer, 2 µl of 1.2 µM RNA, 2 µl of 1 µM TyrRS dimer solution, and 1 µl of ^3^H-labeled L-tyrosine (40 – 60 Ci/mmol). The final concentrations of RNA and TyrRS dimer in the reaction were 120 and 100 nM, respectively. The reactions were incubated for 5 minutes at 30°C. After this time, 80 µl of wash buffer (20 mM Bis-tris pH 6.5, 10 mM NaCl, 1 mM MgCl_2_, and trace amounts of bromophenol blue dye) were added to the reaction, mixed, and quickly filtered through the blotting apparatus described above. An additional 500 µl of washing buffer were flown through the filter. The Hybond membrane was allowed to dry overnight, and the locations marked by the bromophenol blue were cut out. Radioactivity of the Hybond membrane was measured with a scintillation counter as described above.

### Bioinformatic analysis of TLS^Tyr^ variants

To build a sequence and secondary structure consensus model of TLS^Tyr^, we used a preliminary alignment of the TLS^Tyr^ class obtained from the Rfam database (family RF01084) as a starting point, which contained eight sequences lacking the 5’ end of the structure corresponding to stems E and B3 stems (*63, 64*). These eight sequences were manually expanded to include the additional base pairing elements based on published secondary structure models for BMV TLS. Homology searches were performed using Infernal (*65*) to query a database of positive sense RNA viral sequences, resulting in 512 unique sequences derived from eleven distinct viruses, all in the *Bromoviridae* family (Supplemental Files 1 & 2). We evaluated the resulting alignment for nucleotide sequence conservation, as well as significant covariation of base pairing elements, using RNA Structural Covariation Above Phylogenetic Expectation (R-scape) (*66*) analysis to generate a consensus sequence and secondary structure model of TLS^Tyr^ RNAs.

### SHAPE chemical probing

A published protocol was used to probe the secondary structure of representative TLS^Tyr^ variants using *N*-methyl isatoic anhydride (NMIA) as a chemical modifier (*67*). Briefly, the RNAs were folded to their native conformations using the steps described above and reacted for 30 minutes at 23°C with either NMIA or DMSO (control). The reactions contained 3 mg/ml NMIA or DMSO, 50 mM HEPES-KOH pH 8.0, 10 mM MgCl_2_, 60 mM RNA, and 3 nM fluorescently labeled DNA primer for reverse transcription. The reactions were then quenched by adding NaCl to a final concentration of 500 mM and Na-MES pH 6.0 to final concentration of 50 mM. Magnetic beads with poly-dT strands complementary to a poly-A sequence contained within the DNA primer were used to purify the RNA using a magnetic stand (Invitrogen Poly(A)Purist MAG Kit). After purification, the RNAs were resuspended in water to ∼ 240 nM. The RNAs were then reverse transcribed using SuperScript III (Invitrogen) by incubating at 50°C for 45 minutes and degraded by the addition of NaOH to a final concentration of 200 mM followed by incubation for 5 minutes at 90°C. The reactions were quenched by addition of an acidic solution (final concentrations: 250 mM sodium acetate pH 5.2, 250 mM HCl, 500 mM NaCl). To produce RNA ladders, separate reverse transcriptions were performed with each of four ddNTPs (TriLink BioTechnologies). The remaining DNA was then purified using the magnetic stand and eluted in a solution containing HiDi formamide (ThermoFisher) and trace amounts of GeneScan 500 ROX Dye Size Standard (ThermoFisher). Capillary electrophoresis (Applied Biosystems 3500 XL) was used to probe the remaining fluorescently labeled DNA products. Data analysis, including background subtraction and signal normalization using the reactivity of flanking hairpins common to all constructs, was performed using the HiTrace RiboKit computational toolkit (https://ribokit.github.io/HiTRACE/), as described previously (*68–70*).

### Cryo-EM grid preparation and imaging

#### Imaging of BMV TLS RNA

Cryo-EM grids with RNA samples were prepared using a standard procedure. Briefly, 3 µl of folded RNA solution (final concentrations: 10 – 25 µM RNA, 10 mM MgCl_2_, 30 mM Na^+^, 50 mM MOPS pH 7.0) was deposited on C-Flat holey carbon grids (hole size: 1.2 µm; spacing: 1.3 µm; mesh: 400, (VWR)) and a FEI Vitrobot Mark IV was used to plunge freeze the grids. Our initial studies were performed using a ThermoFisher Talos Arctica transmission electron microscope (TEM) operating at an accelerating voltage of 200 kV and equipped with a Gatan K3 Summit direct electron detector (DED). Leginon software was used to collect the data. For wild type BMV TLS RNA, we collected 2389 movies (50 frames) at a magnification of 45,000 (raw pixel size: 0.899 Å) with defocus ranging from –1 to –2.5 µm. The total dose was ∼60 electrons/Å^2^. For BMV_TLS_B3ext_Cshort_ we collected 2494 movies (60 frames) with a total dose of ∼75 electrons/Å^2^ and for BMV_TLS_Dext_B2short_ we collected 505 movies (60 frames) with a total dose of ∼75 electrons/Å^2^. Magnification and defocus range were the same for all constructs. To obtain a higher-resolution map of BMV TLS we used a ThermoFisher Krios TEM equipped with a Falcon 3 DED and a Bioquantum K3 imaging filter at the Pacific Northwest Cryo-EM Center (PNCC). We collected 4463 movies (28 frames) with a total dose of 30 electrons/Å^2^, raw pixel size of 0.415 Å, and a defocus range of –1 to –2.5 µm.

#### Imaging BMV TLS-TyrRS complex

Folded BMV TLS RNA was mixed with purified TyrRS and incubated for 5 minutes at room temperature, then the sample was placed in ice prior to being deposited on C-Flat holey carbon grids and plunge freezing with the Vitrobot. The final solution contained ∼5 uM TyrRS dimer, ∼10 uM BMV TLS RNA, 5 mM MgCl_2_, 250 mM NaCl, 25 mM MOPS (with 15 mM Na^+^ from NaOH titration), 10 mM Tris-HCl, 1 mM β-mercaptoethanol, and 5% glycerol. Preliminary data was collected with the 200 kV Talos Arctica TEM using the same settings as those described above for BMV TLS RNA. Higher-resolution data was collected at PNCC with the 300 kV ThermoFisher Krios. Data was collected at different tilt angles. In total we collected 4302 movies at 0°, 4158 at 20°, and 2502 at 45°. Data at 0° and at 20° were collected from the same grid and data at 45° was collected on a different grid. Movies contained 60 frames and were collected using a pixel size of 0.415 Å, a total dose of 60 electrons/Å^2^, and a defocus range of –1 to –2.5 µm.

#### Imaging of isolated TyrRS homodimer

Three microliters of TyrRS solution containing 10 mM TyrRS dimer, 20 mM Tris-HCl pH 7.5, 500 mM NaCl, 2 mM β-mercaptoethanol, and 10% glycerol was deposited on C-Flat holey carbon grids before plunge freezing with Vitrobot. A dataset of 961 movies (60 frames) was collected with our 200 kV Arctica using the same settings as described above for BMV TLS RNA.

### Cryo-EM data analysis

#### General protocol for all samples

Cryo-EM data was processed using cryoSPARC (*71*). The protocols used for each sample are summarized in Fig. S2-S3 for BMV TLS RNAs, Fig. S11 for the BMV TLS-TyrRS complex, and Fig. S13 for the isolated TyrRS enzyme. For all datasets, movies were imported into cryoSPARC, and motion correction (‘Patch motion correction’ in cryoSPARC) and CTF estimation (‘Patch CTF estimation’ in cryoSPARC) were performed using default parameters. Exposures were curated to eliminate those with obvious defects such as damaged grid areas or excessive ice contamination. Several different methods of picking particles were used, all of which produced essentially identical 3D maps after classification and removal of junk particles, with only small differences in the final resolution of the maps. These methods included manual particle-picking (typically picked > 3000 particles), automated ‘blob picker’ in cryoSPARC, and using low pass filtered templates generated from atomic models. The initial particles picked (by one of the methods described above) were 2D classified to generate templates for automated picking. We also tested Relion (*72*) for the task of generating 2D templates for particle picking, but the quality of the 2D classes was worse than with cryoSPARC. Automatically picked particles were 2D classified to remove ‘junk’ and used for *ab initio* 3D reconstructions using stochastic gradient descent implemented in cryoSPARC (*71*). In some cases, *ab initio* reconstructions were performed without prior 2D classification of the particles, and junk was removed afterwards using 3D classification. These two methods essentially produced the same results, with only minor differences in the final resolution of the maps. 3D classification (heterogenous refinement in cryoSPARC), homogeneous, and non-uniform refinements were performed to produce final maps. Resolution was estimated using gold standard Fourier shell correlation (GSFSC) of 0.143. The resolution reported is corrected for high-resolution noise.

#### Generating 3D maps of BMV TLS RNAs

For BMV TLS RNA with the 200 kV TEM, we manually picked ∼ 3000 particles and classified them to generate templates for automated particle picking (Fig. S2). The putative particles (n = 729, 843) were extracted with a box size of 340 pixels (pixel size: 0.899 Å) and subjected to several rounds of 2D classification to remove junk. The remaining particles (n = 422,021) were used in several *ab initio* 3D reconstructions, varying in the numbers of required classes (Fig. S2). The reconstructions consistently generated a volume with clear RNA secondary structure features (i.e., A-form helices and grooves) and a size consistent with BMV TLS RNA, and other volumes that resembled the main volume but were less defined (Fig. S2). Later atomic modelling of the secondary structure of BMV TLS into the well-defined volume as well as cryo-EM data of BMV TLS RNA bound to TyrRS confirmed that this volume corresponded to the correct folded functional conformation of BMV TLS. To remove potential remaining junk, we generated three *ab initio* volumes and selected the particles that grouped with the well-defined BMV TLS volume (n = 185,596). These particles were used to refine the volume to 7.01 Å resolution (Fig. S2). For BMV TLS RNA with the 300 kV TEM (Fig. S3), we used blob picker on a subset of 500 micrographs and subjected the blobs to 2D classification to generate templates for automated picking in all 9,746 micrographs (Fig. S3). All picked particles were extracted with a 600 pixels box (pixel size = 0.415 Å) and were used to produce three *ab initio* volumes (Fig. S3). The results were consistent with the 200 kV data, reproducing the well-defined volume with RNA secondary structure features and other volumes that were not well defined and were presumably built partly from junk. To remove junk particles, the ‘bad’ volumes were used as particle ‘sinks’ in multiple rounds of 3D classification (‘Heterogenous refinement’ in cryoSPARC) (Fig. S3). The global features of the well-defined BMV TLS RNA volume remained unchanged during each round of 3D classification but the resolution was incrementally better. The particles remaining after several rounds of 3D classification (n = 128,266) were used in homogenous and non-uniform refinements to generate a final map with a resolution of 4.3 Å. Maps for the engineered BMV TLS constructs with extensions and truncations were generated using this same protocol. Stem B3 in modified construct B3^ext^+C^short^, which was 5 bp longer than the original stem, displayed great flexibility and was difficult to visualize in the refined map. To better visualize this stem, we performed a ‘3D Variability Analysis’ in cryoSPARC (*71*) to divide a map into an ensemble and this analysis better revealed the extended B3. In this analysis, the number of modes to solve was set to 3, symmetry to C1 (default), filter resolution to 10 Å, filter order to 1.5 (default), high pass order to 1.5 (default), number of iterations to 20 (default), and Lamda to 0.01 (default). The results from the 3D variability analysis were imported into a ‘3D variability display’ with the output mode set to ‘simple’ and the number of frames set to 20. The first mode was imported into Chimera and displayed as a series of 20 volumes, some of which more clearly showed the position of the extended B3. Importantly, none of the other helices (except for the shortened C) displayed noticeable differences relative to wild type BMV TLS. The volume with the best defined B3 was used to fit a structural model (Fig. S7). A similar analysis was performed with wild type BMV TLS to generate Fig. 1F, with filter resolution set to 7.5 and number of frames set to 4 (as shown in Fig. 1F). The four volumes generated from this analysis were then used to reclassify the particles and each volume was refined separately using ‘homogeneous refinement’ in cryoSPARC.

#### Generating 3D maps of TLS-TyrRS complex

For the complex, we started our analysis with micrographs collected without tilt (n = 4302). To generate initial 2D templates, we picked particles on a subset of 126 micrographs using ‘blob picker’ in cryoSPARC with particle size range set to 70–200 Å (Fig. S11A). The templates were used for automated particle picking in the 4302 micrographs. After removal of junk particles using 2D classification, 374,993 particles were used to generate *ab initio* 3D reconstructions that clearly showed density consistent with BMV TLS RNA bound to TyrRS (Fig. S11A). Only one bound RNA appeared well defined in the reconstructions, as explained in Results and Discussion. A model generated from these initial reconstructions was used to simulate 2D templates for picking particles in all the micrographs, including those with 0°, 20°, and 45° tilts (Fig. S11B). We then performed several rounds of 2D classification to remove junk particles and combined the particles from all tilts, resulting in 783,472 particles that were used for the *ab initio* reconstruction of four maps (Fig. S11B). The maps clearly showed BMV TLS RNA bound to TyrRS in two distinct conformations. Several rounds of 3D classification followed by homogeneous and non-uniform refinements generated maps with 5.5 and 6.0 Å resolution maps for each of the two conformations (Fig. S12).

#### Generating 3D map of isolated TyrRS dimer

We used the crystal structure of yeast TyrRS (PDB ID: 2DLC) to simulate a low-resolution (10 Å) EM map using Chimera (*73*) and produce 2D templates for particle picking. The particles were extracted with a 300 pixels box (pixel size: 0.899 Å). After one round of 2D classification to remove junk, we separated 2D projections that corresponded to the ‘long’ dimensions of TyrRS from those that corresponded to the ‘short’ dimensions, as done previously for an elongated protein of similar dimensions (*74*). Particles in the ‘short’ classes (n = 708,269) were subjected to a second round of classification using a circular mask of 120 Å and ‘bad’ classes (e.g., displaying more than one particle) were discarded. After discarding bad classes, 386,859 particles were used for *ab initio* reconstruction of three models, one of which showed dimensions consistent with TyrRS (Fig. S13). This map was refined to 9.19 Å resolution using non-uniform refinement in cryoSPARC.

### Structural modeling

#### Modeling of unbound BMV TLS RNA using molecular dynamics flexible fitting and real-space refinements and autoDRRAFTER

A schematic of the methods used to build the BMV TLS RNA structure is provided in Fig. S6. We began with the helical assignments obtained from the helical extensions/truncations and the published computational model of the BMV TLS RNA (*75*), which agreed well with the known secondary structure. Each helical domain was extracted from the published model and docked into the corresponding density using UCSF Chimera (*73*). Manual modeling was performed in Chimera and Coot (*76*) to join the helical domains and to obtain an initial model for flexible fitting (Fig S6A). We then performed molecular dynamics flexible fitting using the web server Namdinator (https://namdinator.au.dk/namdinator/) (*77*), with the following parameters: Map resolution: 4.3 Å, Start temperature: 298 K, Final temperature: 298 K, G-force Scaling Factor: 0.2, Minimization steps: 4000, Simulation steps: 20,000, Phenix RSR cycles: 1, Implicit solvent (GBIS): exclude. The resulting model was imported into Phenix (*78*) along with the cryo-EM map to perform additional real-space refinements (RSR), maintaining the secondary structure of the model during refinements. ‘Minimization_global’, ‘local_grid_search’ and ‘adp’ were selected in Phenix RSR, with Max Interations: 50, Macro cycles: 1, Target bonds rmsd: 0.0005, and Target angles rmsd: 1.5. Then the model and the cryo-EM map were imported into ERRASER to correct the RNA geometry (*79, 80*). Default settings were used in ERRASER. Several iterative rounds of PHENIX RSR and ERRASER were used to incrementally decrease steric clashes and improve the geometry of the model. Finally, we used ‘Geometry minimization’ under Model Tools in Phenix. The final structural model of BMV TLS displayed a MolProbity clashscore of 8.69, 0 bad bonds, 0 bad angles, 28 outlier backbone conformations, and 0 probably wrong sugar puckers (Table S1). These statistics were obtained using the MolProbity web server (http://molprobity.biochem.duke.edu) (*81*). Additional model statistics and parameters are given in Table S1. The outlier conformations were examined and were generally located in complex areas of the structure expected to have non-canonical conformations. The same method was used to build structural models for engineered constructs B3^ext^+C^short^ and D^ext^+B2^short^ (Fig. S7).

As an orthogonal method for modelling the structure of BMV TLS RNA we used autoDRRAFTER to generate ten models (*82*) (Fig. S6B). To do this we followed the steps described in the Rosetta Commons website (https://new.rosettacommons.org/docs/latest/application_documentation/rna/auto-drrafter#running-auto-drrafter) with the following parameters: convergence_threshold 10, nmodels 1000, njobs 10. The RMSD between the manually built structure and the autoDRRAFTER models were obtained using the ‘align’ command in Pymol (Schrödinger, Inc).

#### Modelling of BMV TLS-TyrRS complex

To generate an initial homology model for *Phaseolus vulgaris* TyrRS monomer we used Modeller (*83*) implemented within UCSF Chimera (*73*). Modeller used the 2.21 Å crystal structure of the *Plasmodium falciparum* TyrRS (template sequence; PDB ID: 3VGJ), which displayed the greatest score in a multiple sequence alignment with our target sequence. A segment of 37 residues was not modeled because there was no homology with the template sequence. The model obtained from Modeller was imported into the I-Tasser web server (*84*) to build the missing 37 residues while keeping the rest of the structure fixed. Two copies of the I-Tasser TyrRS monomer and the structure of unbound BMV TLS RNA were imported into Chimera and docked into the cryo-EM maps of the BMV TLS-TyrRS complex. Domain B3+E was rotated manually into the corresponding density. The resulting model of the complex was then imported into the Namdinator web server (*77*) and MDFF was set with the following parameters: Map resolution: 5.5 or 6.0 Å (bound states 1 and 2, respectively), Start temperature: 298 K, Final temperature: 298 K, G-force Scaling Factor: 0.2, Minimization steps: 4000, Simulation steps: 40,000, Phenix RSR cycles: 1, Implicit solvent (GBIS): exclude. The models were imported into Phenix to perform geometry minimization, followed by addition of hydrogens using the Molprobity web server. Model statistics are provided in Table S2 & S3 for bound states 1 and 2, respectively.

---

**Fig. S1.**
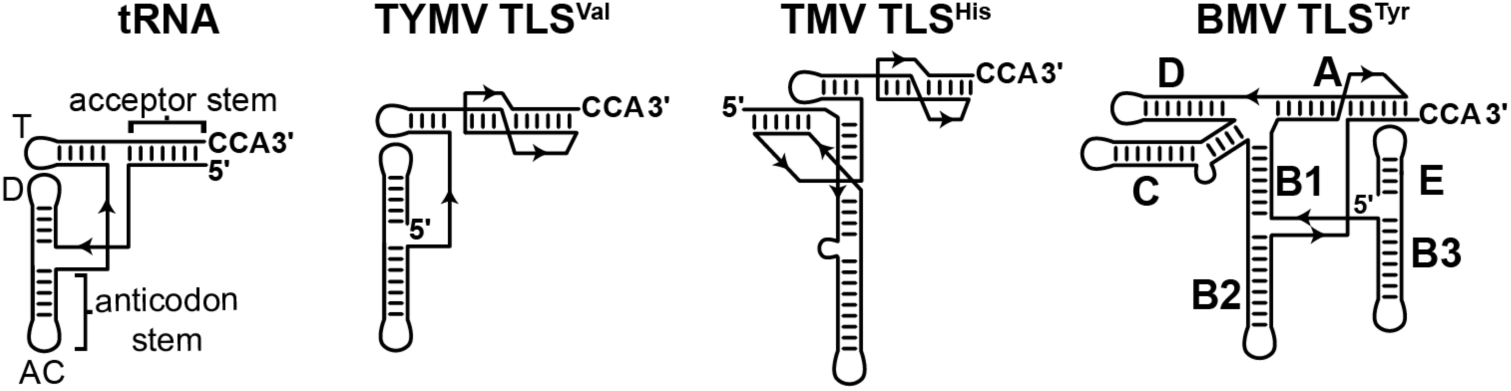
Comparison of the secondary structures of tRNA and TLS from different classes. TLSs are classified based on their aminoacylation specificities. Three classes have been identified in viruses that infect plants: valylatable (TLS^Val^), histidylatable (TLS^His^), and tyrosylatable (TLS^Tyr^) (*85*). Representatives from each class are shown. These are turnip yellow mosaic virus (TYMV), tobacco mosaic virus (TMV), and brome mosaic virus (BMV). The canonical structural features of tRNA are labeled: the anticodon and acceptor stems and the T, D and anticodon (AC) loops. The helical stems of BMV TLS are labeled with the names assigned in previous studies (*75*).

**Fig. S2.**
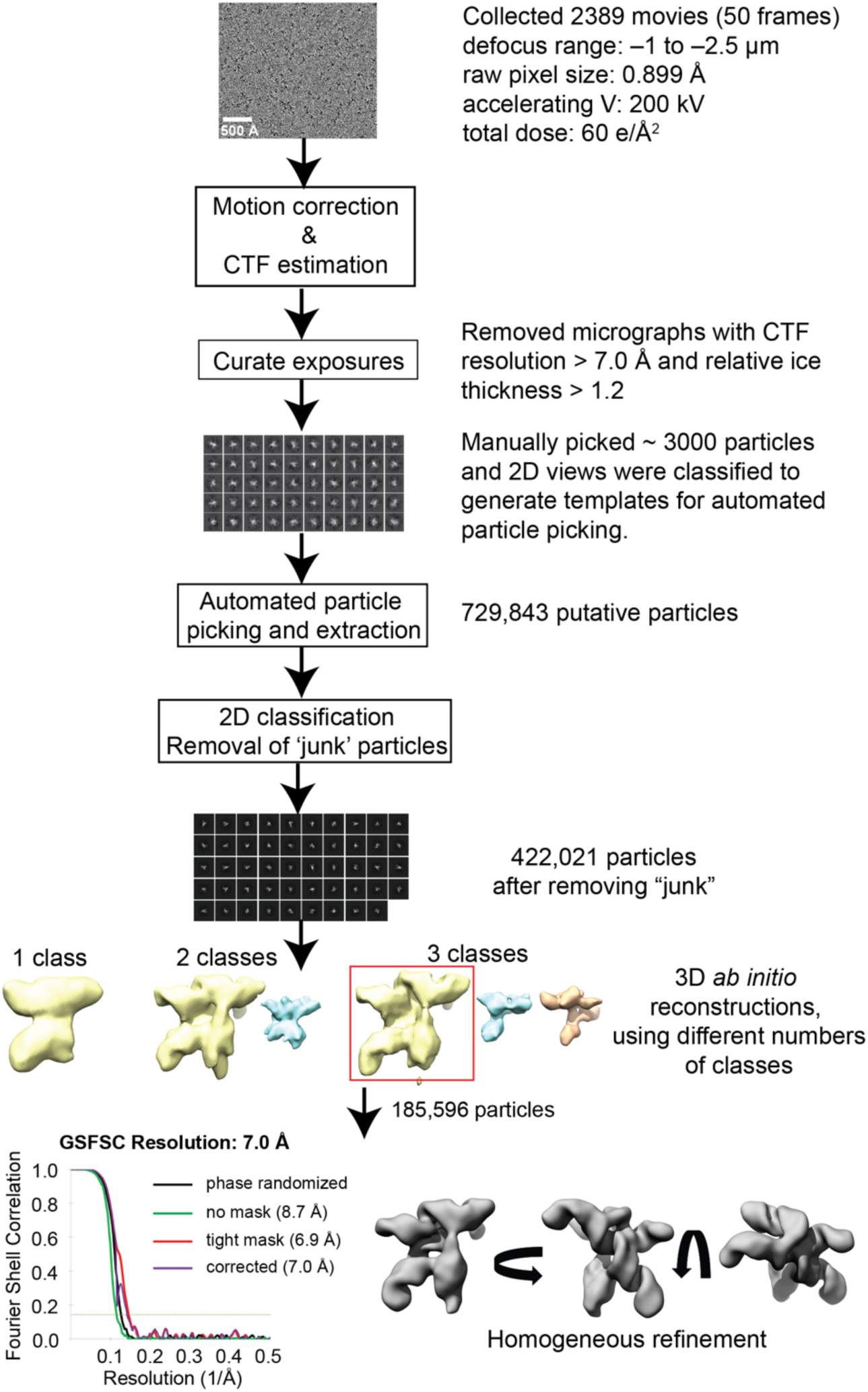
Analysis of BMV TLS RNA data collected with 200 kV electron microscope. Steps followed to generate cryo-EM reconstruction of BMV TLS using cryoSPARC (*71*). *Ab initio* reconstructions were performed using different numbers of possible models. In every case, there was a reconstruction that was consistent with the size and expected topology of BMV TLS and that had clear RNA secondary structural features (yellow volumes). Cryo-EM experiments in the presence of TyrRS (see main text) confirmed that these maps corresponded to the functional conformation of BMV TLS. The BMV TLS map created with three classes was selected as an initial model for further refinement. The overall resolution of the refined map was estimated using half maps and gold standard FSC (GSFSC) of 0.143. The resolution reported (7.01 Å) was corrected using high-resolution noise substitution to measure the amount of noise overfitting in cryoSPARC (*86*).

**Fig. S3.**
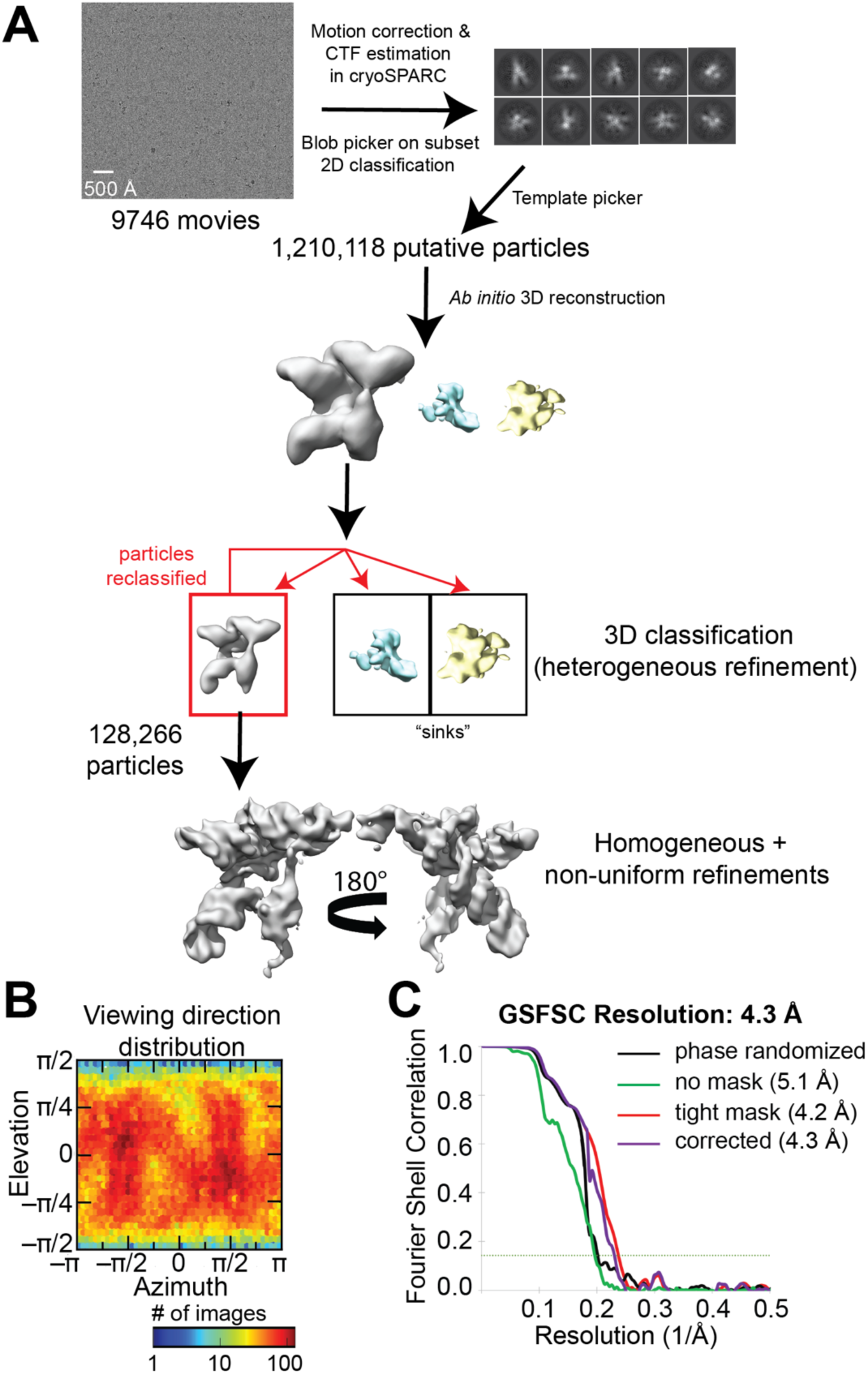
Analysis of BMV TLS RNA data collected with 300 kV electron microscope. (A) Steps followed to generate cryo-EM reconstruction of BMV TLS using cryoSPARC (*71*). After *ab initio* reconstruction, several rounds of 3D classification (Heterogeneous refinement) were performed to increase the resolution of the map, using bad volumes as ‘sinks’. The final map (4.3 Å resolution) was generated using homogeneous refinement followed by a non-uniform refinement with 128,266 particles. (B) Distribution of viewing angles. (C) Estimation of overall resolution of cryo-EM maps using half maps and gold standard FSC of 0.143.

**Fig. S4.**
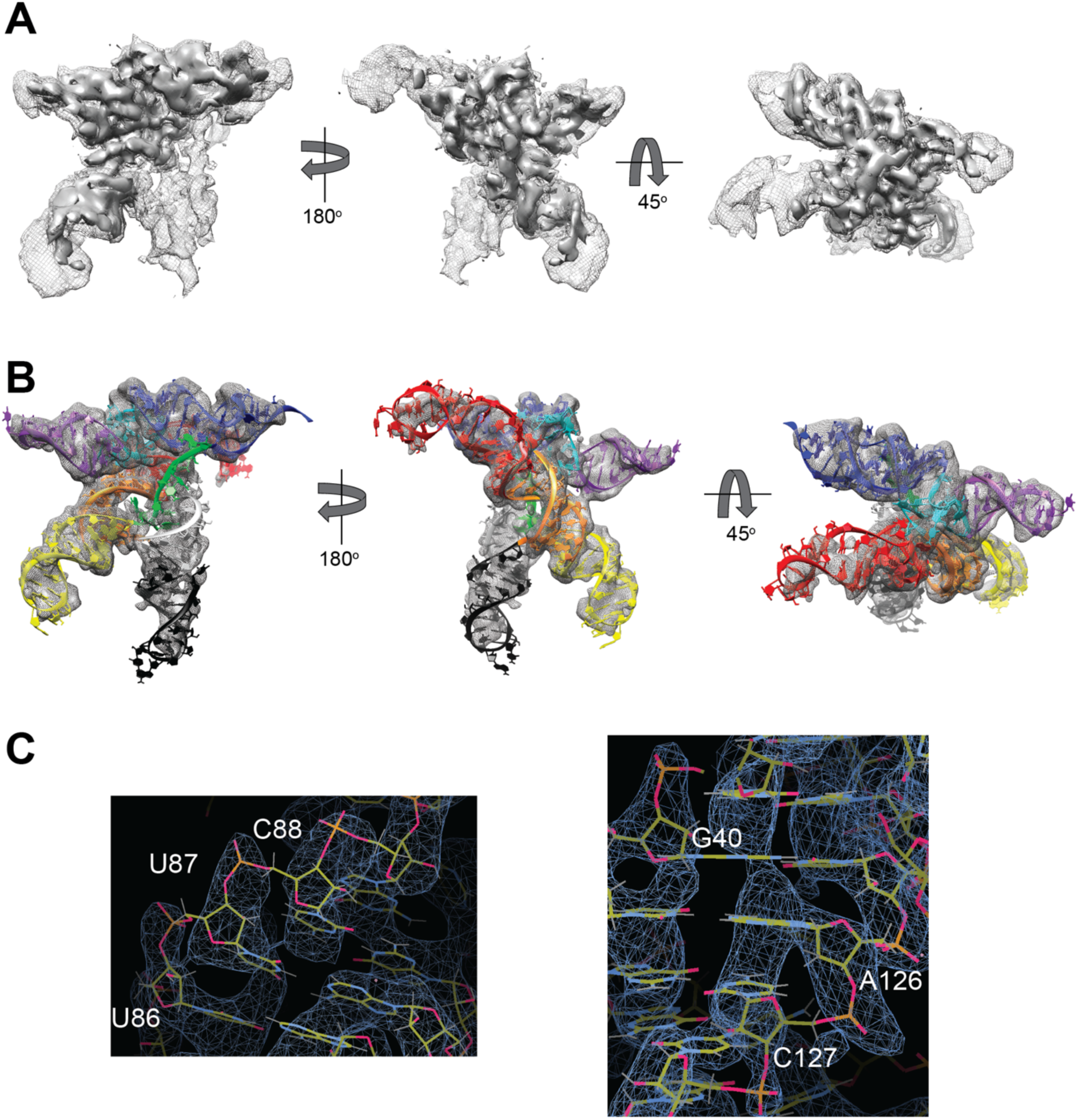
Model fit to density. (A) 4.3 Å resolution map at two different contour levels to display different features of the map. The map was generated using homogeneous refinement followed by a non-uniform refinement in cryoSPARC with 128,266 particles. The reported resolution was corrected for high-resolution noise overfitting using noise substitution and phase randomized maps (*86*). (B) Structural model of BMV TLS fitted to cryo-EM density. Colors are as in Fig. 2 (C) Examples of density displaying phosphate ‘bumps’ and base stacking. Map was sharpened using density modification in Phenix (*87*). Figures were generated in Coot (*76*).

**Fig. S5.**
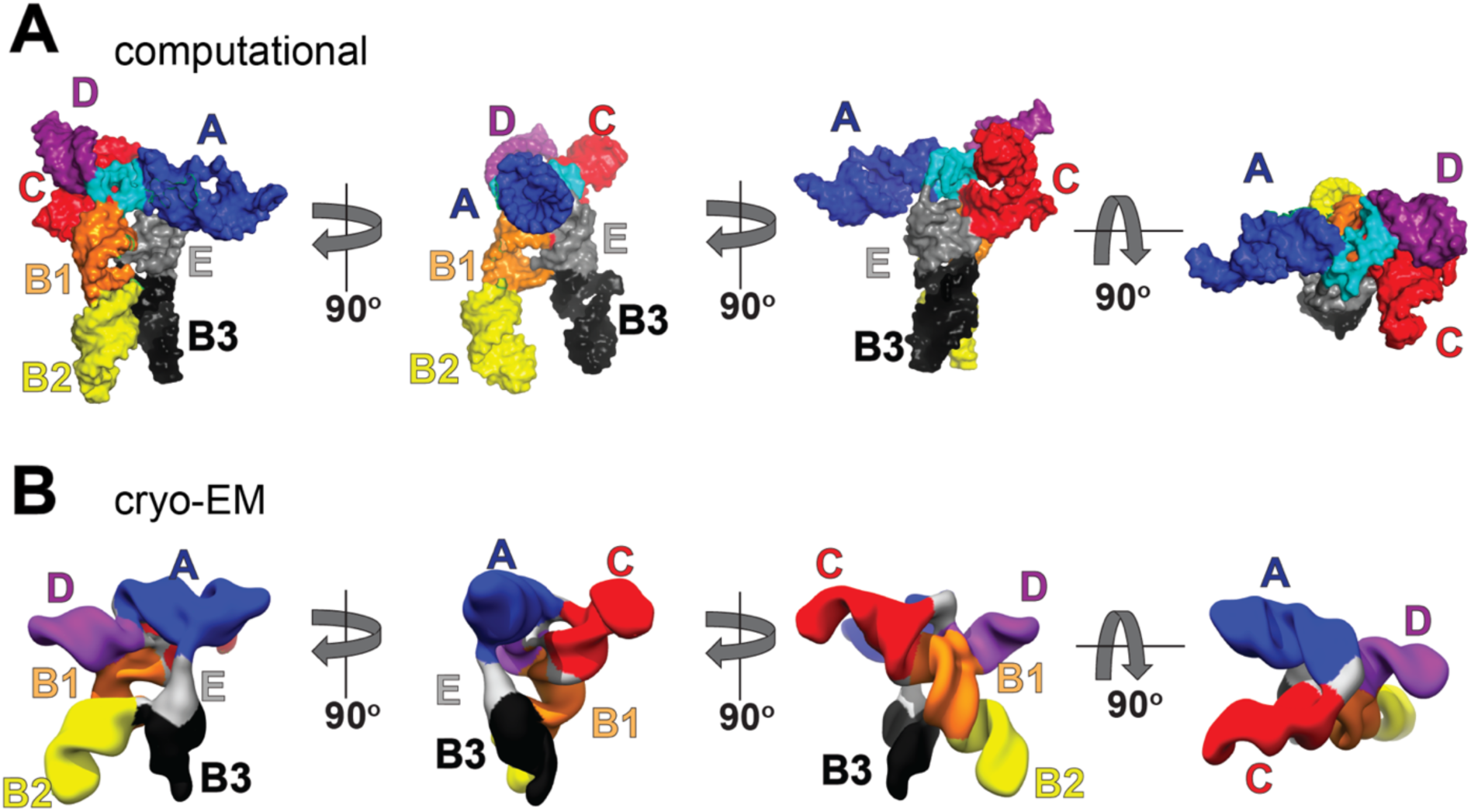
Comparison between cryo-EM map and published computational model. Helices are colored according to secondary structure in Figure 2A of the main text. (A) Previously published computational model from chemical and enzymatic probing data and phylogenetic secondary structure comparisons (*75*). The structure is shown using a solvent-accessible surface representation. (B) The map obtained by cryo-EM is oriented similarly to the computational model in (A) for comparison, with helical elements colored as in panel A. Although global features of the cryo-EM map and the computational model are similar, the location and orientation of the helices differ.

**Fig. S6.**
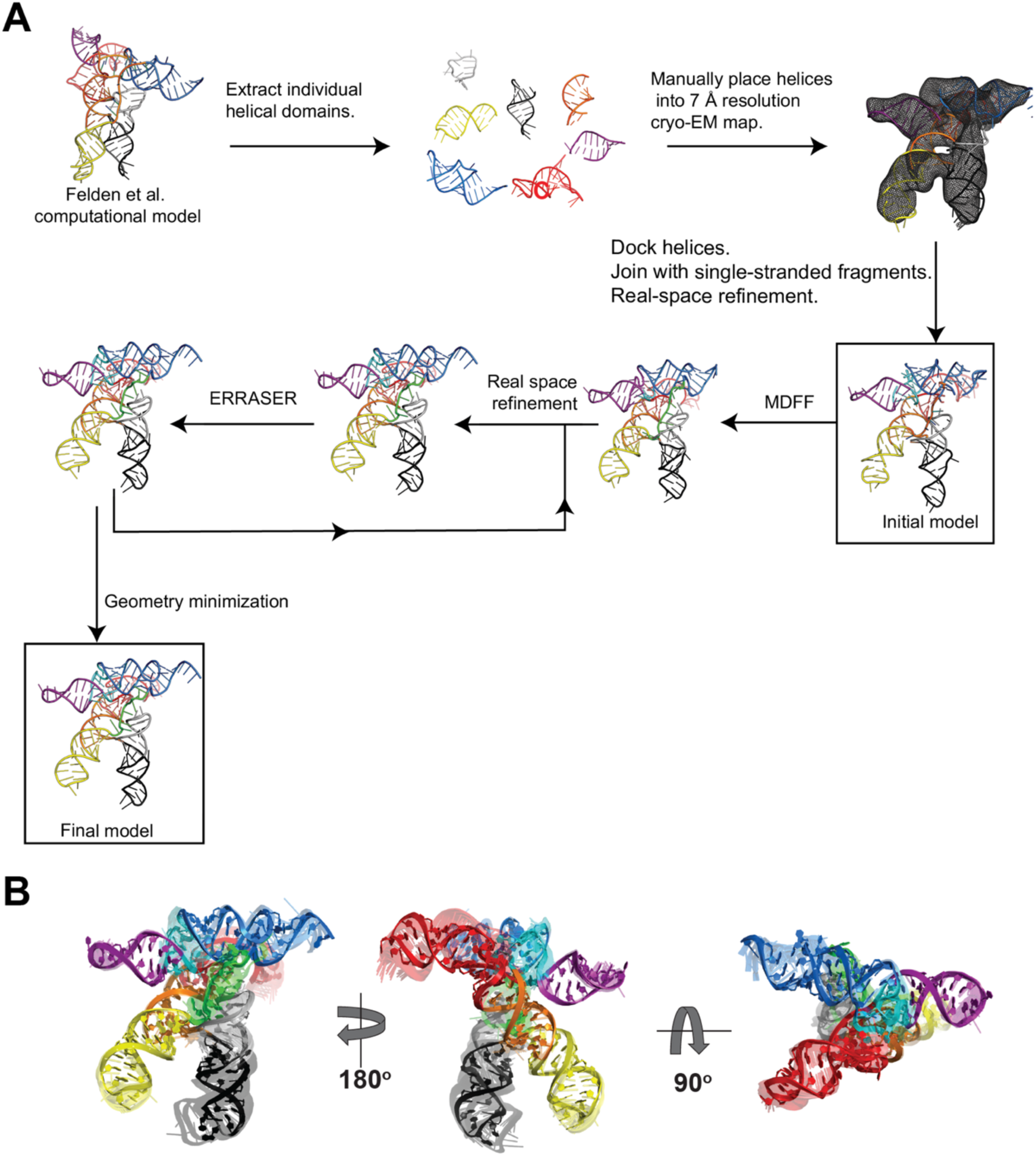
Structural modelling of BMV TLS. (A) Building a structural model of BMV TLS using manual and automated modelling tools. Details of this process, as well as parameters used, are given in the Materials and Methods section. Briefly, a structural model of BMV TLS was built starting from the computational model of Felden et al., as this model is largely consistent with the experimentally determined secondary structure (*75*). Individual helices were extracted from the computational model and docked into a moderate resolution (∼7Å) cryo-EM map using the ‘fit in map’ function in UCSF Chimera (*73*). Single-stranded regions connecting the helices were added using Chimera and Coot (*76*). A real space refinement (RSR) in Phenix (*78*) was performed to build an initial model of BMV TLS that fit into the low-resolution map. Then we used Namdinator (*77*) to perform molecular dynamics flexible fitting (MDFF; see Material and Methods for critical parameters), using a 4.3 Å map. The model obtained from the MDFF job was then imported back into Phenix for an additional RSR, using the secondary structure of BMV TLS as a constraint. To correct the geometry of the RNA we used ERRASER (*79, 80*). Several iterations of Phenix RSR and ERRASER were performed to lower the MolProbity clash score and improve the geometry of the model (http://molprobity.biochem.duke.edu) (*81*). A ‘Geometry minimization’ in Phenix was performed at the end. The final model displayed a MolProbity clashscore of 8.69, 0 bad bonds, 0 bad angles, 28 outlier backbone conformations, and 0 probably wrong sugar puckers (Table S1). (B) Computational modelling of BMV TLS using autoDRRAFTER. Ten computational models were generated using autoDRRAFTER (transparent/colored) and are superimposed over the structural model of BMV TLS (opaque/shaded) generated by the method described in (A). autoDRRAFTER is part of the Ribosolve pipeline for accelerated generation of structural models of RNA-only molecules using cryo-EM maps and takes a cryo-EM map, a sequence file, and a secondary structure model as inputs (*82*). The computational models agreed well with our structural model, with RMSD values ranging from 2.0 to 3.2 Å.

**Fig. S7.**
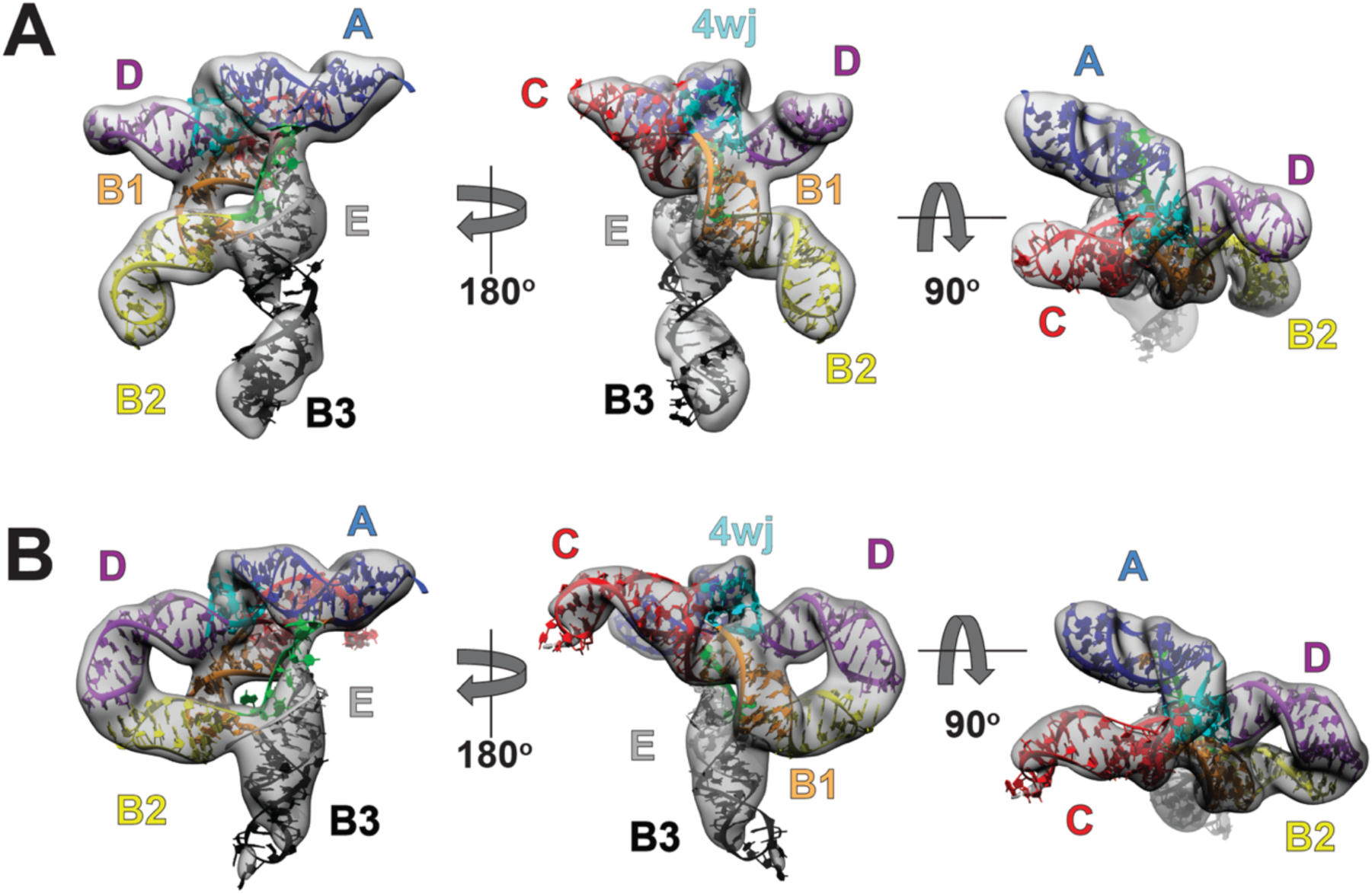
Structural models of modified constructs support architecture of BMV TLS. (A) Density of B3^ext^+C^short^, with the structural model docked inside. In this model, extra base pairs have been added to stem B3 and base pairs have been removed from stem C. The density matches a model with these altered helical lengths. (B) Same as panel A, but with D^ext^+B2^short^. The lengthening of stem D and the shortening of B2 in the model match the density. Unexpectedly, these changes induced a new tertiary contact between the apical loops of B2 and D, apparently due to a fortuitous alignment of the bases in their apical loops.

**Fig. S8.**
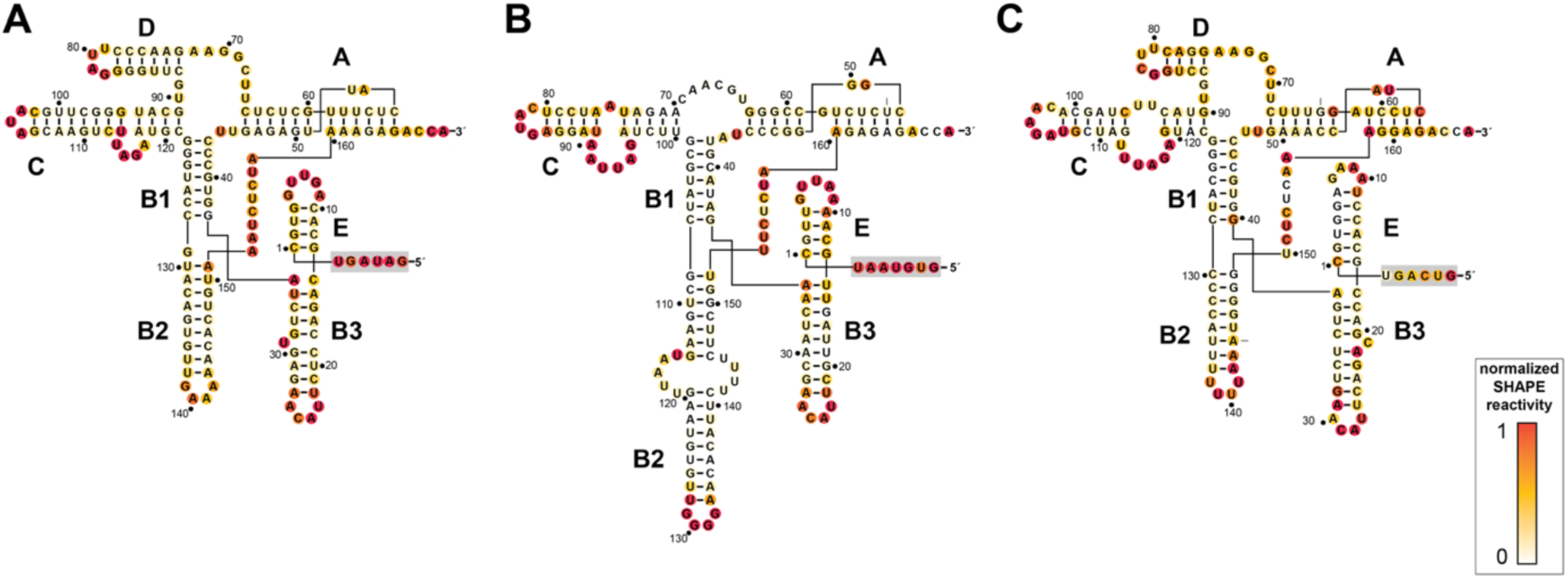
Chemical probing of representative TLS^Tyr^ variants supports secondary structure consensus model. SHAPE probing of TLS^Tyr^ belonging to three different viruses: (A) BMV, (B) pelargonium zonate spot virus, and (C) cucumber mosaic virus. Helices were labeled using the BMV TLS naming. Structures were drawn to match the consensus sequence and secondary structure model in the main text (Figure 3). Variant from pelargonium zonate spot virus lacks a stem D. 5’ sequences in grey box were added for chemical probing and are not part of the model.

**Fig. S9.**
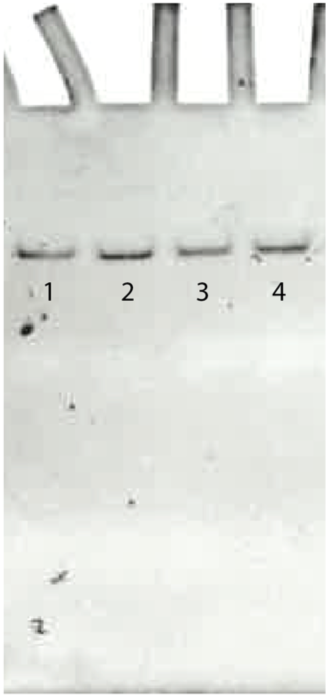
Gel electrophoresis under native condition of BMV TLS RNA constructs in which specific apical loops were mutated to UUCG. (1) Wild type BMV TLS RNA, (2) E (UUCG), (3) B2 (UUCG), and (4) B3 (UUCG) were folded and run in a 10% polyacrylamide gel with 10 mM MgCl_2_. The running buffer contained 33 mM TRIS-HCl, 64 mM HEPES, pH 7.4, and 10 mM MgCl_2_. Electrophoresis was performed in a cold room at 4°C. RNA was stained with ethidium bromide and imaged with a UV scanner.

**Fig. S10.**
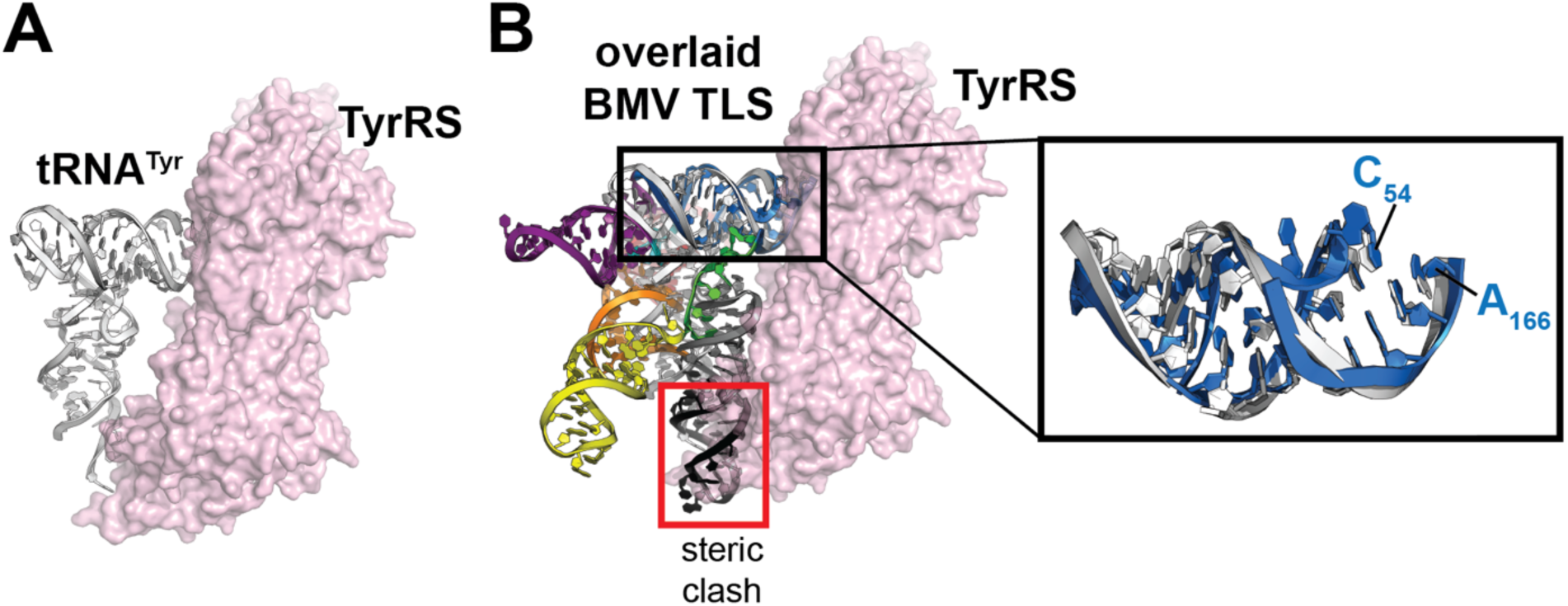
Modelling of BMV TLS bound to TyrRS suggests that conformational changes are required. (A) Published structure of yeast tRNA^Tyr^ bound to TyrRS homodimer (*88*). Two tRNAs bind to each TyrRS homodimer; for simplicity only one tRNA is shown. (B) Identical structure to panel (A), with BMV TLS superimposed on tRNA^Tyr^ by aligning the structures of the acceptor stems. Boxed: Close-up of the aligned acceptor stems, with tRNA in grey and the BMV TLS stem A in blue. The alignment was performed prioritizing base pairs near the terminal CCA, as this region is known to be critical for tyrosylation (*88, 89*). Selected nucleotides are labeled for reference.

**Fig. S11.**
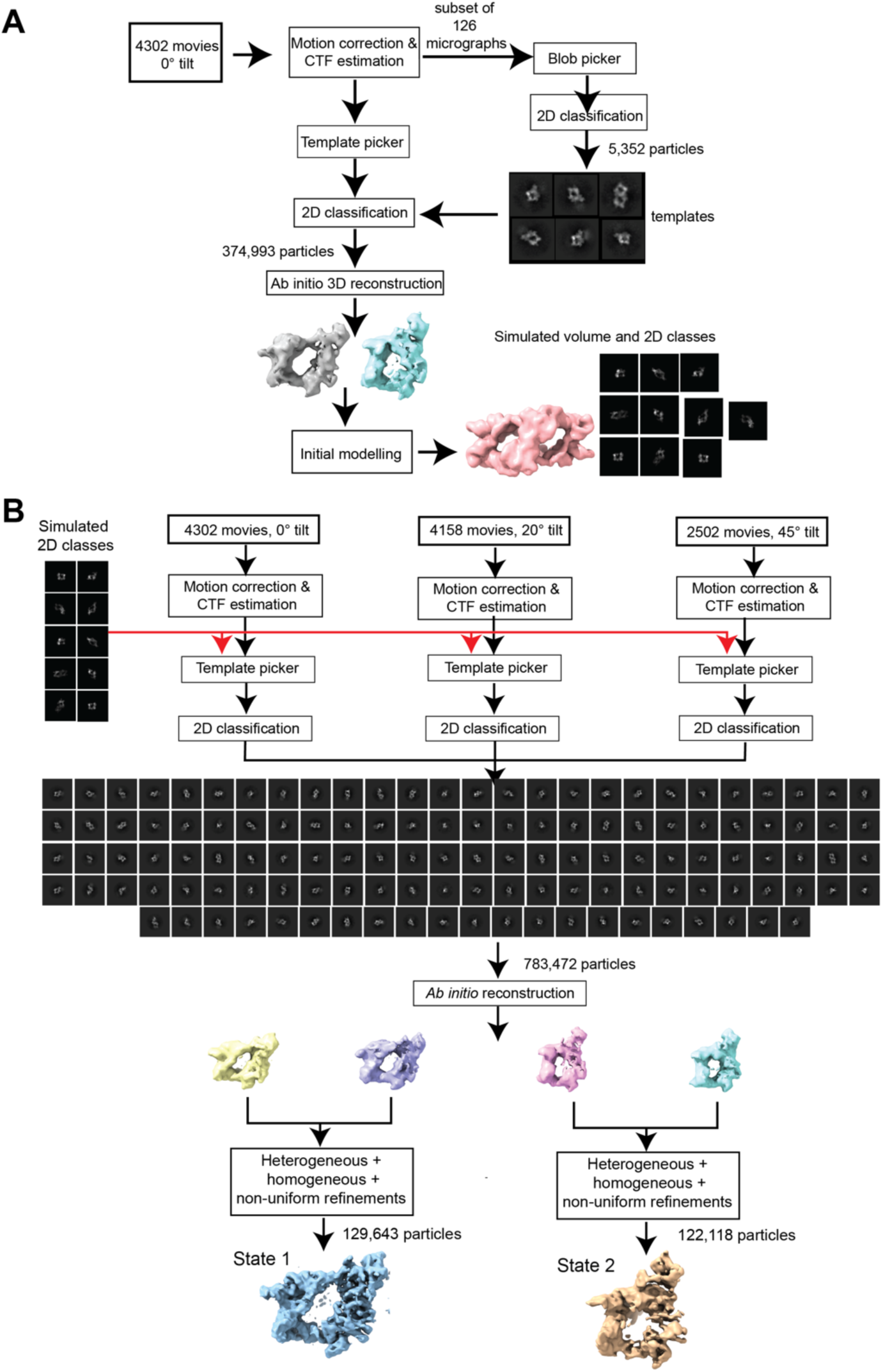
Analysis of cryo-EM data of the BMV TLS-TyrRS complex. (A) Preliminary analysis of the cryo-EM data collected without tilt. Templates for automated particle picking were generated using blob picker on a subset of 126 micrographs. *Ab initio* reconstructions showed density consistent with BMV TLS RNA bound to TyrRS. These reconstructions were used to generate an initial atomic model of the full complex and a set of simulated 2D projections. (B) Analysis using data collected at three different angles (0°, 20°, and 45°) to reduce preferred orientations. Templates generated in (A) were used to pick particles in all micrographs. The simulated 2D templates were low pass filtered before particle picking, so details of the model do not bias the picking. Two distinct conformations of the bound complex were observed. Although two bound RNAs are seen in multiple 2D classes, one RNA is well-defined in the 3D reconstructions. Bound state 1 was refined to 5.5 Å and bound state 2 to 6.0 Å (Fig. S12).

**Fig. S12.**
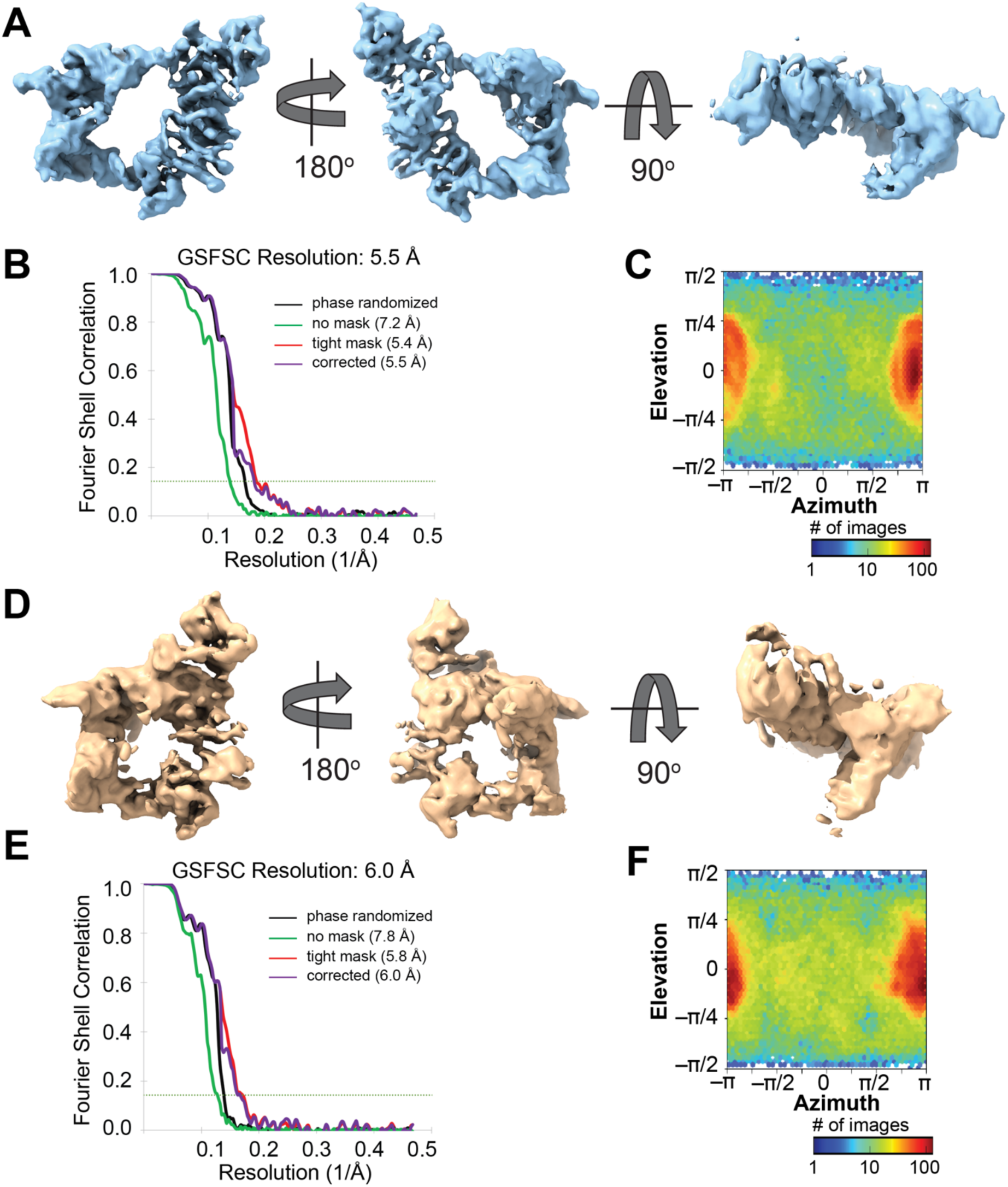
Refined volumes of two bounds states of the BMV TLS-TyrRS complex. (A) Cryo-EM map of bound state 1, viewed in three different orientations. (B) FSC curves for refined map of bound state 1. Resolutions reported are calculated using gold standard FSC (GSFSC) of 0.143. The ‘Corrected’ resolution was corrected for high resolution noise using phase randomized maps. (C) Distribution of views in the refined map of bound state 1. (D) Cryo-EM map of bound state 2, viewed in three different orientations. (E) FSC curves for refined map of bound state 2. Resolutions reported are calculated using gold standard FSC (GSFSC) of 0.143. The ‘Corrected’ resolution was corrected for high resolution noise using phase randomized maps. (F) Distribution of views in the refined map of bound state 2. Above the graphs in B and E, put a space in between the number and the A symbol.

**Fig. S13.**
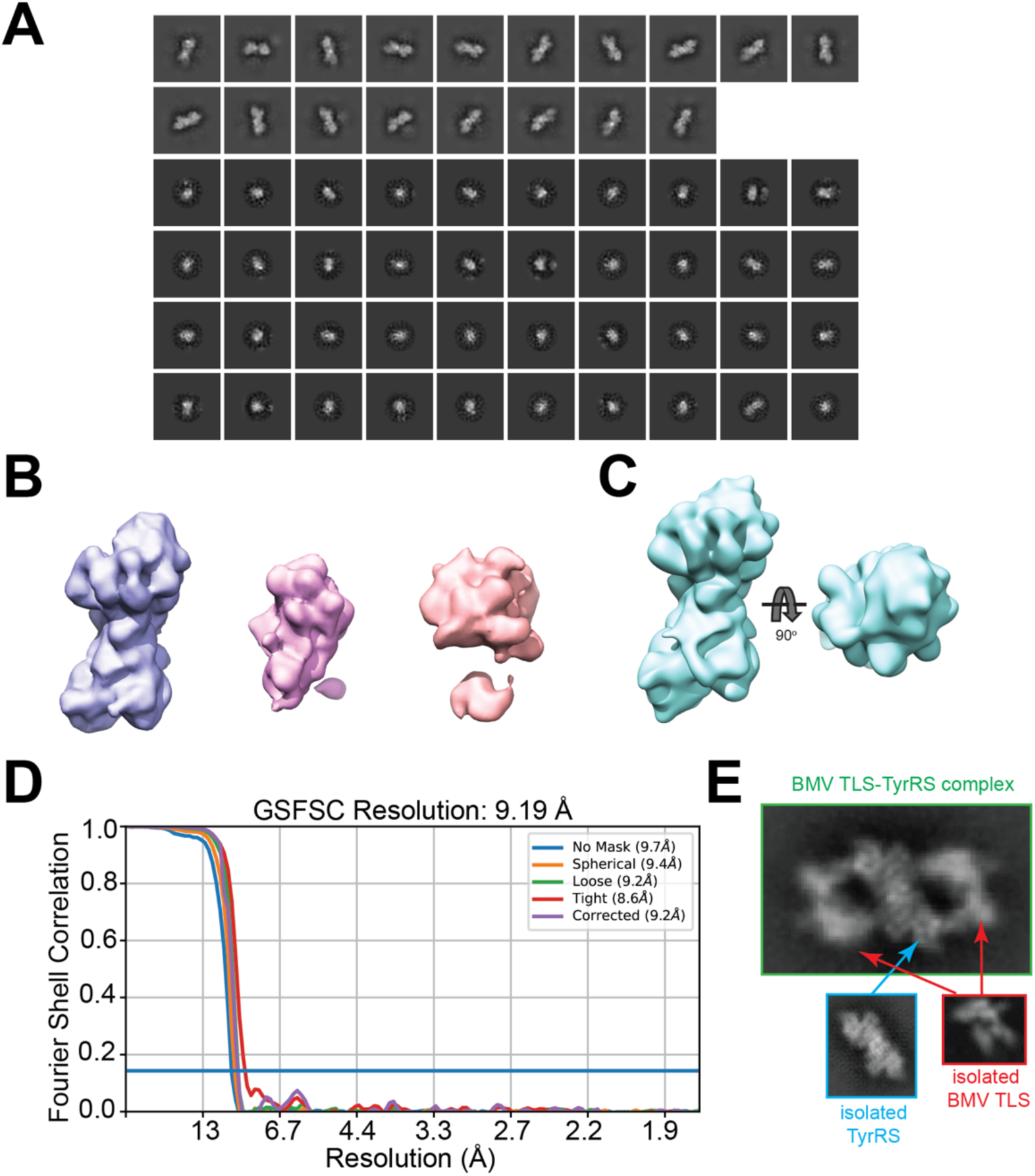
Cryo-EM studies of purified TyrRS from *Phaseolus vulgaris.* (A) 2D classes of TyrRS after removal of junk particles via 2D classification and selection. Classification showed two major types of classes: side views (long classes; first two rows) and top views (short classes; four bottom rows). Because of the large differences in the dimensions of these two types of classes, they were separated and classified using circular masks of different sizes, as done previously for a protein of similar dimensions (*74*). (B) *Ab initio* 3D reconstructions generated using stochastic gradient descent with cryoSPARC (*71*). The reconstruction on the left (purple volume) agreed with the dimensions expected for TyrRS. (C) Refined volume of TyrRS. (D) Fourier shell correlation (FSC) curves. The overall resolution of the refined map was estimated using half maps and gold standard FSC of 0.143. The resolution reported (9.19 Å) was corrected using high-resolution noise substitution to measure the amount of noise overfitting; the correction utilizes phase randomized maps (*86*). (E) Comparisons of a 2D class of BMV TLS-TyrRS complex (green box), isolated TyrRS (cyan box), and isolated BMV TLS RNA (red box). Density in the middle of the complex corresponds to TyrRS (cyan arrow). Two molecules of BMV TLS RNA can be observed bound to TyrRS (red arrows).

**Fig. S14.**
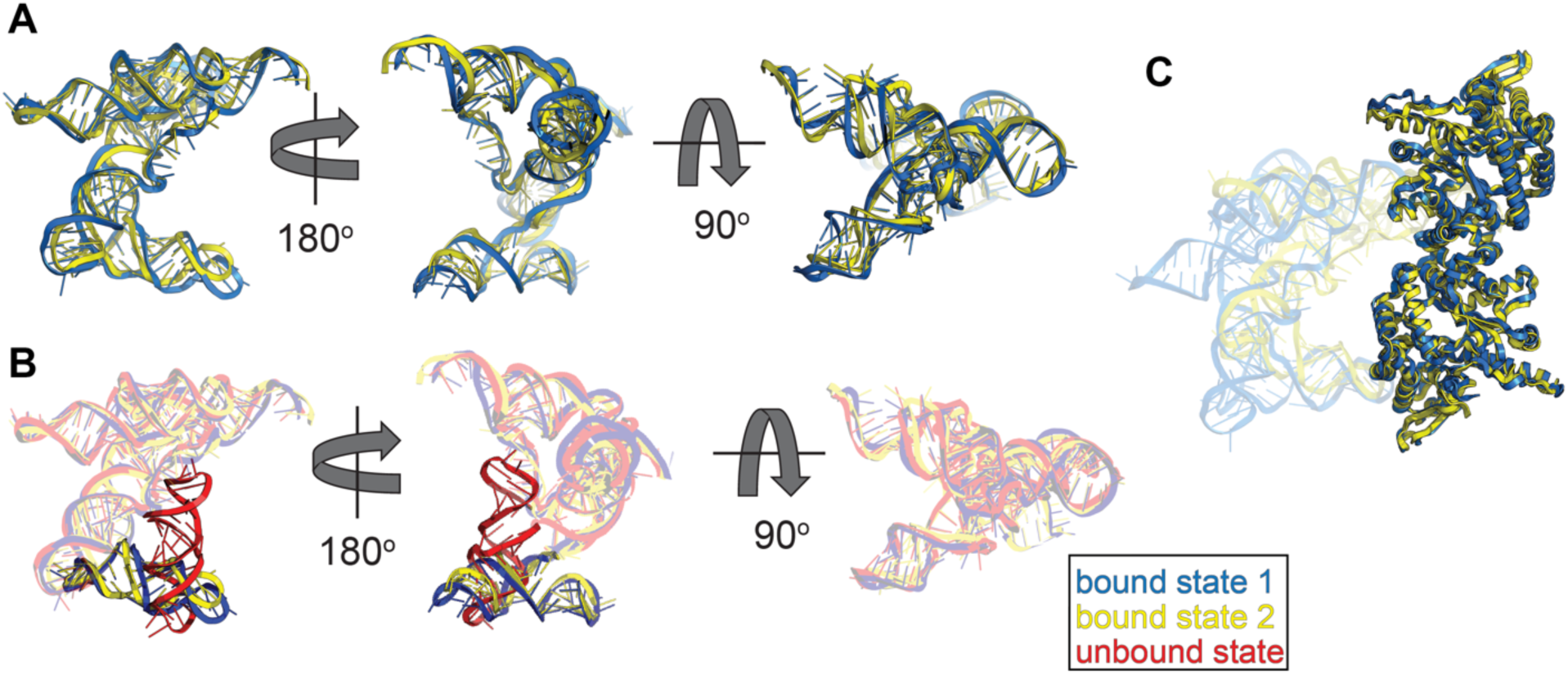
Comparison between bound and unbound states of BMV TLS RNA. (A) BMV TLS RNA adopts the same global conformation in both bound states. Analysis of the cryo-EM data of BMV TLS-TyrRS complex revealed two different conformational states: bound state 1 (blue) and bound state 2 (yellow). In bound state 1, the acceptor arm of the RNA is not fully docked into the active site (see Fig. 4 in main text). In bound state 2, the acceptor arm is deeper into TyrRS and the 3’ CCA is positioned near the active site. Despite these differences, the global conformation of the BMV TLS RNA is essentially the same in both bound states, with an RMSD of 2.66 Å as calculated using the ‘align’ command in Pymol. (B) Superposition of bound and unbound BMV TLS RNA reveals large conformational changes. In the unbound state (red), B3+E on average occupies a position at roughly a right angle to the acceptor stem analog. In that position B3+E would clash with the TyrRS. In the bound states 1 and 2 (blue and yellow, respectively) B3+E has rotated approximately 90 degrees relative to the unbound conformation and lays roughly parallel to the acceptor stem analog, avoiding any steric clash with the enzyme and placing the B3 apical loop anticodon analog on the surface of TyrRS. (C) The conformation of the TyrRS is essentially the same in both bound states but its position relative to BMV TLS RNA differs. Structures of bound states 1 and 2 are superimposed based on the alignment of TyrRS. The RMSD between the proteins is 1.6 Å as calculated using the ‘align’ command in Pymol. The bound BMV TLS RNA is transparent.

**Table S1.**
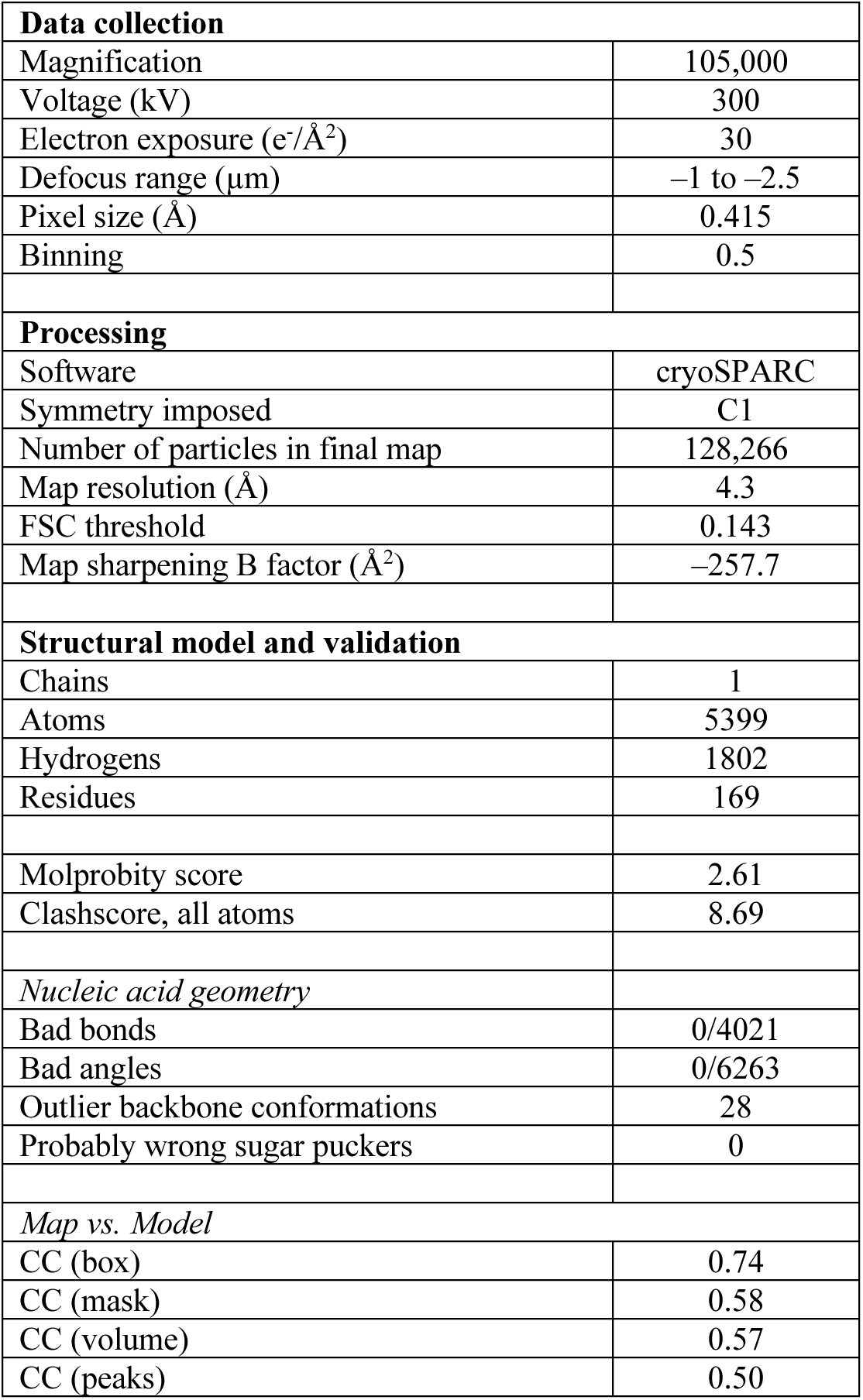
Cryo-EM data collection parameters, characteristics of final map, and validation of the structural model of BMV TLS RNA. Statistics were obtained using ‘Comprehensive validation (cryo-EM)’ in Phenix (*78*) and the MolProbity web server (http://molprobity.biochem.duke.edu) (*81*).

**Table S2.**
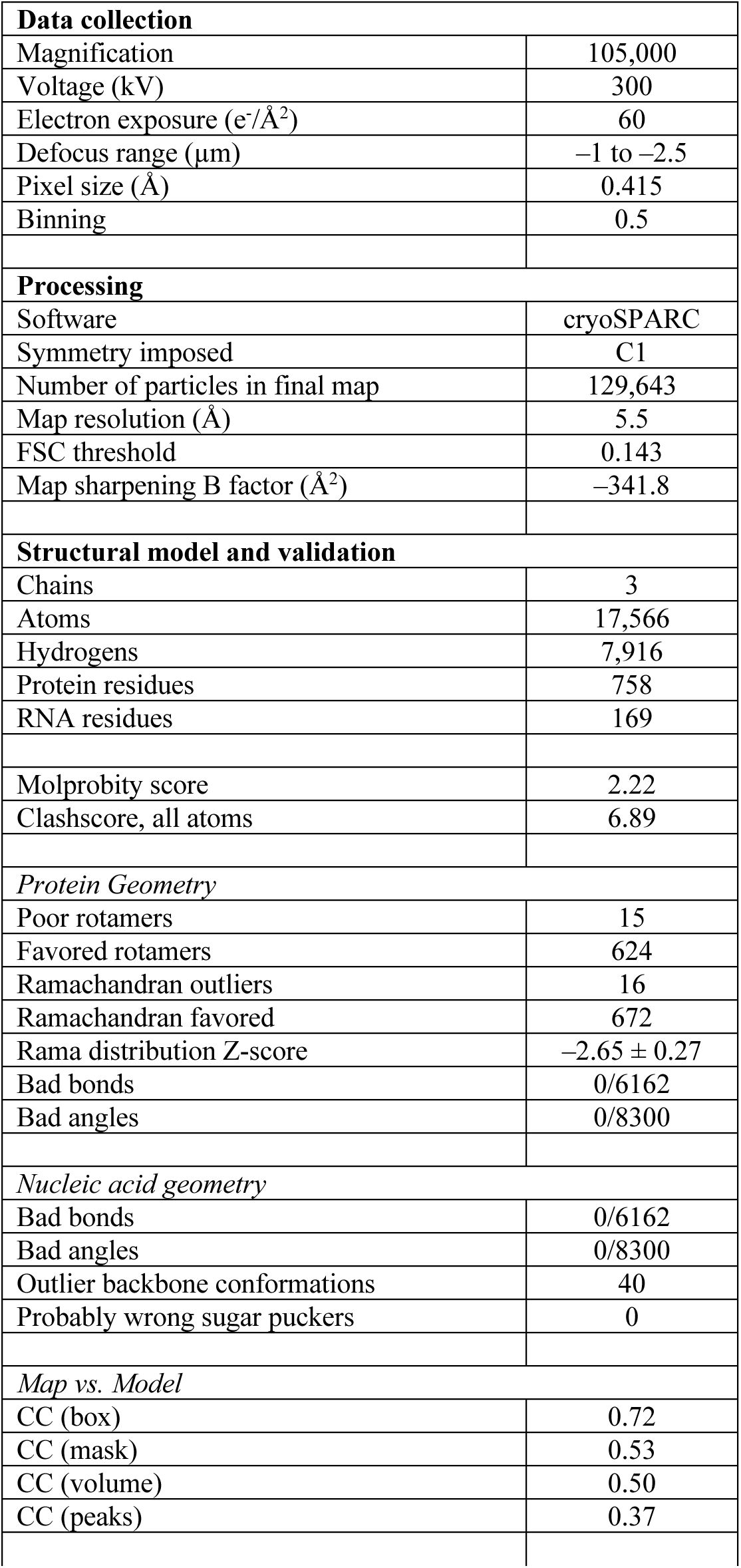
Cryo-EM data collection parameters, characteristics of final map, and validation of the structural model of the BMV TLS-TyrRS complex in bound state 1. Statistics were obtained using ‘Comprehensive validation (cryo-EM)’ in Phenix (*78*) and the MolProbity web server (http://molprobity.biochem.duke.edu) (*81*).

|  |  |
| --- | --- |
| **Data collection** |  |
| Magnification | 105,000 |
| Voltage (kV) | 300 |
| Electron exposure (e<sup>-</sup>/Å<sup>2</sup>) | 60 |
| Defocus range (μm) | -1 to -2.5 |
| Pixel size (Å) | 0.415 |
| Binning | 0.5 |
| **Processing** |  |
| Software | cryoSPARC |
| Symmetry imposed | C1 |
| Number of particles in final map | 129,643 |
| Map resolution (Å) | 5.5 |
| FSC threshold | 0.143 |
| Map sharpening B factor (Å<sup>2</sup>) | -341.8 |
| **Structural model and validation** |  |
| Chains | 3 |
| Atoms | 17,566 |
| Hydrogens | 7,916 |
| Protein residues | 758 |
| RNA residues | 169 |
| Molprobity score | 2.22 |
| Clashscore, all atoms | 6.89 |
| *Protein Geometry* |  |
| Poor rotamers | 15 |
| Favored rotamers | 624 |
| Ramachandran outliers | 16 |
| Ramachandran favored | 672 |
| Rama distribution Z-score | -2.65 ± 0.27 |
| Bad bonds | 0/6162 |
| Bad angles | 0/8300 |
| *Nucleic acid geometry* |  |
| Bad bonds | 0/6162 |
| Bad angles | 0/8300 |
| Outlier backbone conformations | 40 |
| Probably wrong sugar puckers | 0 |
| *Map vs. Model* |  |
| CC (box) | 0.72 |
| CC (mask) | 0.53 |
| CC (volume) | 0.50 |
| CC (peaks) | 0.37 |

**Table S3.** Cryo-EM data collection parameters, characteristics of final map, and validation of the structural model of the BMV TLS-TyrRS complex in bound state 2. Statistics were obtained using ‘Comprehensive validation (cryo-EM)’ in Phenix (*78*) and the MolProbity web server (http://molprobity.biochem.duke.edu) (*81*).

|  |  |
| --- | --- |
| **Data collection** |  |
| Magnification | 105,000 |
| Voltage (kV) | 300 |
| Electron exposure (e<sup>-</sup>/Å<sup>2</sup>) | 60 |
| Defocus range (μm) | -1 to -2.5 |
| Pixel size (Å) | 0.415 |
| Binning | 0.5 |
| **Processing** |  |
| Software | cryoSPARC |
| Symmetry imposed | C1 |
| Number of particles in final map | 122,118 |
| Map resolution (Å) | 6.0 |
| FSC threshold | 0.143 |
| Map sharpening B factor (Å<sup>2</sup>) | -398.4 |
| **Structural model and validation** |  |
| Chains | 3 |
| Atoms | 17,566 |
| Hydrogens | 7,916 |
| Protein residues | 758 |
| RNA residues | 169 |
| Molprobity score | 2.28 |
| Clashscore, all atoms | 10.25 |
| *Protein Geometry* |  |
| Poor rotamers | 10 |
| Favored rotamers | 633 |
| Ramachandran outliers | 18 |
| Ramachandran favored | 658 |
| Rama distribution Z-score | -3.26 ± 0.27 |
| Bad bonds | 6/6162 |
| Bad angles | 0/8300 |
| *Nucleic acid geometry* |  |
| Bad bonds | 0/6162 |
| Bad angles | 1/8300 |
| Outlier backbone conformations | 43 |
| Probably wrong sugar puckers | 3 |
| *Map vs. Model* |  |
| CC (box) | 0.71 |
| CC (mask) | 0.47 |
| CC (volume) | 0.41 |
| CC (peaks) | 0.30 |

**Table S4.**
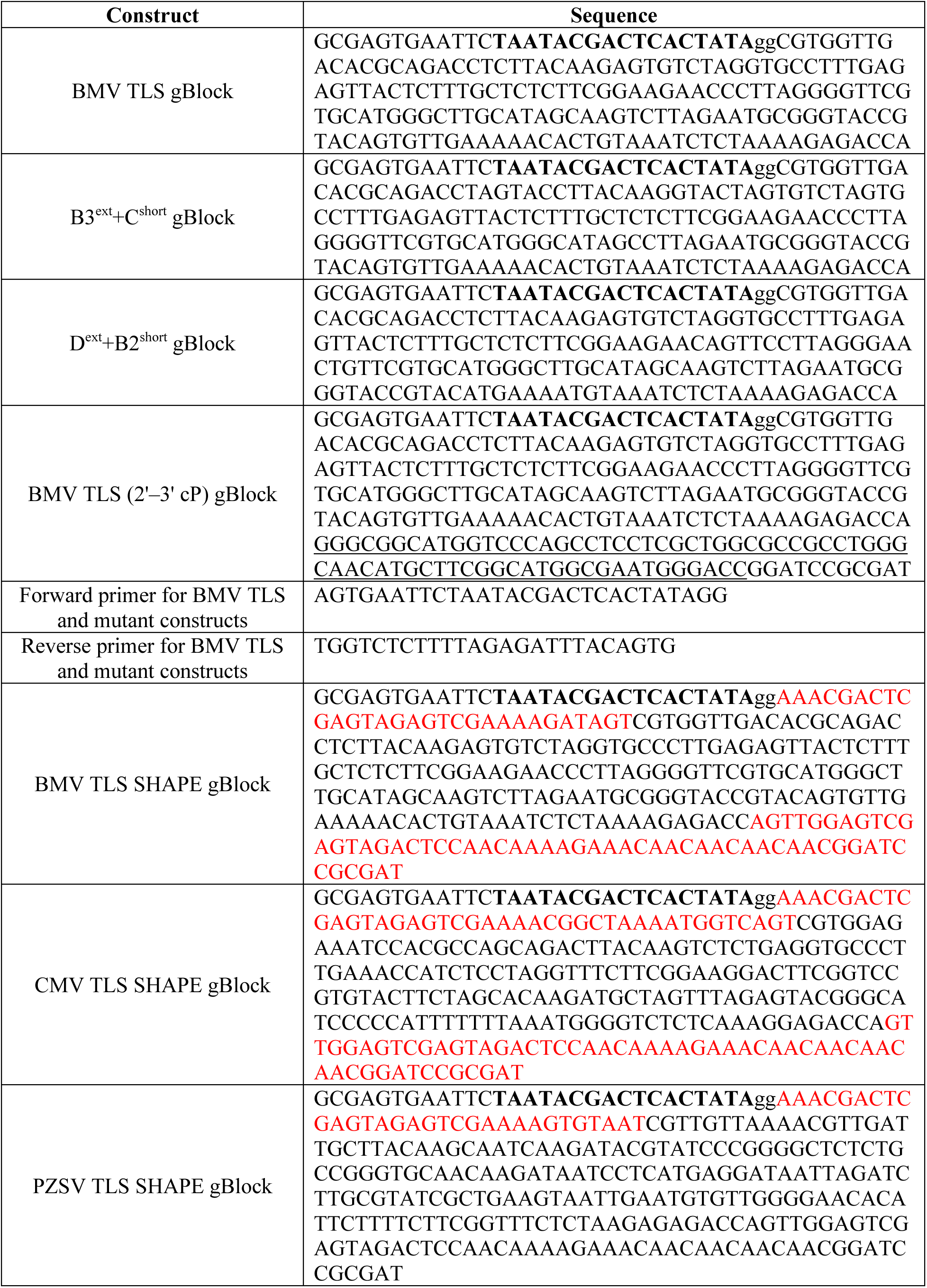
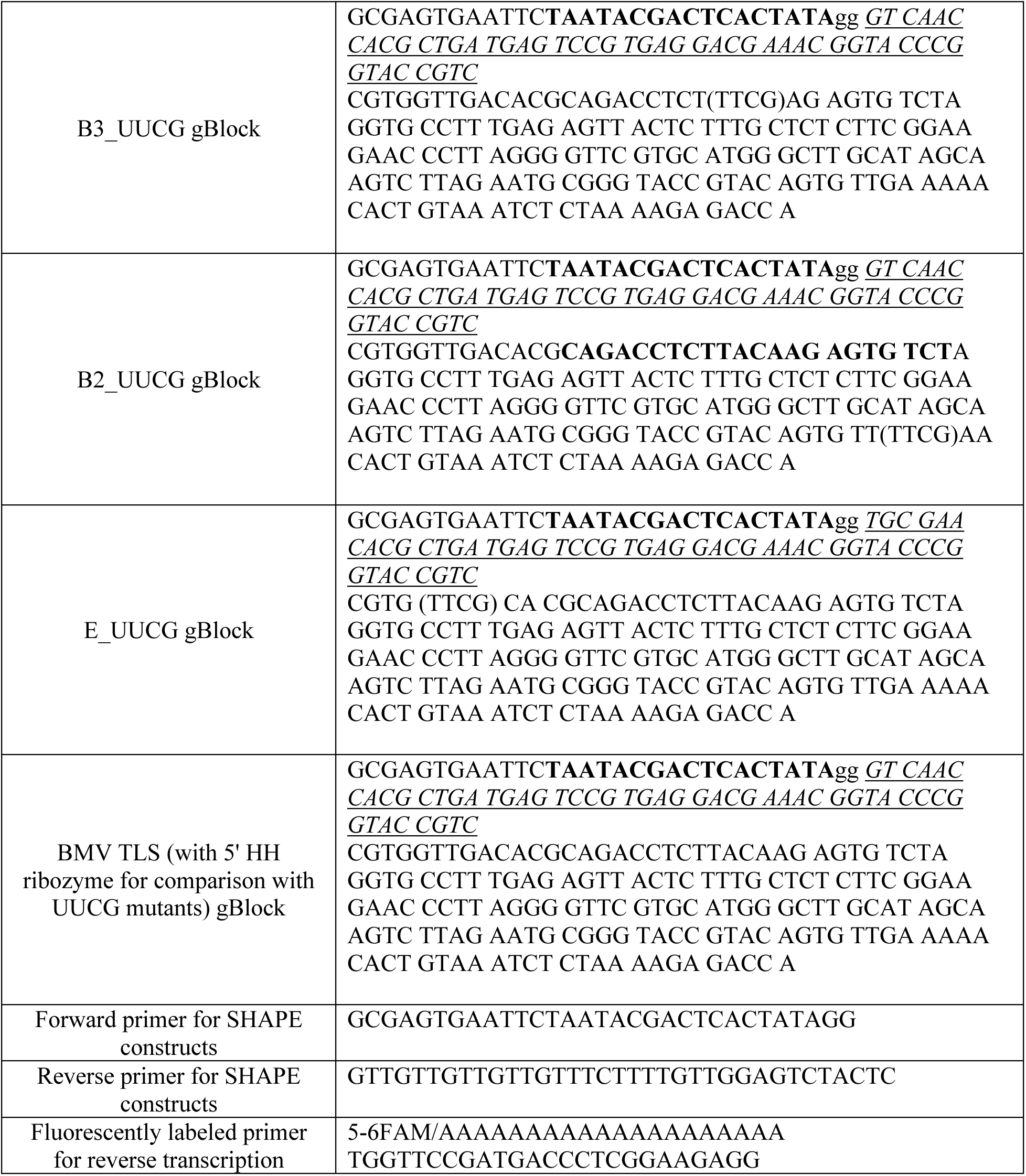
Sequences of constructs used. Sequence in bold is T7 promoter. Lower case ‘gg’ was added to facilitate *in vitro* transcription. Sequence in regular font and underlined corresponds to hepatitis delta virus (HDV) ribozyme at the 3’ end of desired sequence. Sequence in italic and underlined corresponds to hammerhead (HH) ribozyme at the 5’ end of desired sequence. Ribozymes are cleaved as described in Materials and Methods. Sequences in red were added for SHAPE experiments and contain hairpins used for signal normalization (ref. 53). Sequences in parenthesis are UUCG mutations. All sequences were ordered from Integrated DNA Technologies.

## Notes

### Competing Interest Statement

The authors have declared no competing interest.

### Summary of Updates

In the first version, we reported how the structure of the BMV TLS RNA, solved by cryo-EM, was highly suggestive of conformational dynamics in the unbound form and of structural changes upon synthetase binding. However, the nature of these changes and the RNA-protein complex was speculative. We have now solved the structure of BMV TLS RNA bound to a plant tyrosyl synthetase. This structure and the resultant conclusions are described in this revised manuscript, as are additional 3D Variability Analysis, the use of automated building programs, and in vitro aminoacylation assays. Briefly, by comparing the structures of the bound and unbound forms of the RNA, we can now directly observe conformational rearrangements inherent in one domain of the unbound RNA and describe the nature of these rearrangements. This domain contains the anticodon mimic, and it undergoes a substantial structural shift with synthetase binding. The bound state differs dramatically from that adopted by tRNA.

